# Efficient *in vivo* pharmacological inhibition of ΔFOSB, an AP1 transcription factor, in brain

**DOI:** 10.1101/2025.10.21.683721

**Authors:** Sean McNeme, Anil Kumar, Yun Young Yim, Brandon W. Hughes, Corey St. Romain, Yi Li, Ashwani Kumar, Qichao Bao, Molly Estill, Shanghua Fan, Nadeen Takatka, Earnest P. Chen, Matthew Rivera, Haiying Chen, Alfred J. Robison, Mischa Machius, Steve J. Haggarty, Jeannie Chin, Eric J. Nestler, Jia Zhou, Gabby Rudenko

## Abstract

ΔFOSB, an unusually stable member of the AP1 family of transcription factors, mediates long-term maladaptations that play a key role in the pathogenesis of drug addiction, cognitive decline, dyskinesias, and several other chronic neurological and psychiatric conditions. We have recently identified that 2-phenoxybenzenesulfonic acid-containing compounds disrupt the binding of ΔFOSB to DNA *in vitro* in cell-based assays, and one such compound, JPC0661, disrupts ΔFOSB binding to genomic DNA *in vivo* in mouse brain with partial efficiency. JPC0661 binds to a groove outside of the DNA-binding cleft of the ΔFOSB/JUND bZIP heterodimer in the co-crystal structure. Here, we generated a panel of analogs of JPC0661 with the goal of establishing structure-activity relationships and improving its *in vivo* efficacy by replacing the amino-pyrazolone cap moiety with various substituents. We show that one such analog, YL0441, disrupts the binding of ΔFOSB to DNA *in vitro* and *in vivo*, and suppresses ΔFOSB-function in cell-based assays. Importantly, infusion of YL0441 into the hippocampus of APP mice (a mouse model for Alzheimer’s disease) leads to virtually complete loss of ΔFOSB bound to genomic DNA by CUT&RUN sequencing. Our findings corroborate that DNA binding/release of AP1 transcription factors can be controlled via small molecules, even by analogs of a compound that binds to a groove outside of the DNA-binding cleft, and that our lead can be optimized via medicinal chemistry to yield a highly efficacious inhibitor of ΔFOSB function *in vivo*. These findings define a strategy to design small-molecule inhibitors for other AP1- and AP1-related transcription factors.

**IN BRIEF:** We demonstrate the creation of a highly effective inhibitor, YL0441, of ΔFOSB, an AP1 transcription factor, which decreases the number of ΔFOSB-bound sites to genomic DNA by ∼94% upon *in vivo* infusion to the hippocampus of APP mice, a mouse model for Alzheimer’s disease. This work generates a highly novel probe compound to assess the therapeutic value of ΔFOSB *in vivo*, a transcription factor with a critical role in mediating long-term changes in gene expression in several neuropsychiatric disorders in addition to Alzheimer’s disease, including drug addiction, seizure-related cognitive decline, and dyskinesias.

## INTRODUCTION

ΔFOSB, an AP1 transcription factor, has emerged as a promising drug target for neurological and psychiatric disorders, including drug addiction, cognitive decline, and dyskinesias. The ΔFOSB protein accumulates in specific regions of the brain in response to a range of chronic (but not acute) stimuli (Robison & Nestler, 2022). For example, ΔFOSB protein levels rise in the striatum in response to drugs of abuse and mediate increased drug seeking and intake (Teague & Nestler, 2022; Robison & Nestler, 2022). Likewise, ΔFOSB protein levels rise in the striatum in response to antipsychotic medications or to L-DOPA, which are used to treat a variety of psychotic disorders and Parkinson’s disease, and ΔFOSB is thought to play a key role in mediating the debilitating involuntary movements that accompany these treatments (Beck *et al*, 2019, 2021). Very high levels of ΔFOSB protein are also found in the hippocampus of individuals with epilepsy or with Alzheimer’s Disease (AD) examined postmortem, and the magnitude of expression corresponds with the severity of cognitive impairment (You *et al*, 2017; Corbett *et al*, 2017; Fu *et al*, 2023).

The ΔFOSB protein, which is formed through alternative splicing of the *FosB* gene transcript and lacks the C-terminal 101 residues of full-length FOSB, is uniquely stable and has a half-life of ∼7 days *in vivo* in brain while FOSB and all other FOS family proteins are much less stable with a half-life of several hours at most (Ulery *et al*, 2006; Ulery-Reynolds *et al*, 2009; Manning *et al*, 2017; Teague & Nestler, 2022). ΔFOSB stably alters gene expression by binding to AP1 DNA consensus sites (AP1 site; 5’-TGA C/G TCA-3’) in the genome at specific regulatory sites, which mediates a variety of long-term neural and behavioral adaptations (Robison & Nestler, 2022). ΔFOSB binds to promoter regions of select target genes, where it can decrease or increase gene expression, but most ΔFOSB binds to AP1 sites at putative enhancer elements distributed throughout intergenic regions and gene bodies, where it impacts target gene expression from these distal locations (Robison & Nestler, 2022; Yeh *et al*, 2023; McNeme *et al*, 2025). The ΔFOSB protein (237 residues) contains an intrinsically disordered N-terminal region (∼150 a.a.), which likely recruits various cofactors, while the C-terminal basic region/leucine zipper (bZIP) forms a forceps-shaped molecule upon assembly with another AP1 bZIP partner that generates a DNA-binding cleft located at the fulcrum of the forceps (Jorissen *et al*, 2007; Yin *et al*, 2017, 2020). In brain, the most common dimerization partner of ΔFOSB is thought to be the AP1 transcription factor JUND (Chen *et al*, 1995; Hiroi *et al*, 1998). Dimerization of ΔFOSB and its partner via their respective leucine zippers enables the DNA-binding motifs (a helical basic region in the bZIP domain) to insert into the major groove of DNA (Jorissen *et al*, 2007; Yin *et al*, 2017). DNA binding is further controlled by two cysteine residues found N-terminal in the DNA-binding motifs in the bZIP domains (ΔFOSB Cys^172^ and JUND Cys^285^ in human) that form a redox switch (Yin *et al*, 2017; Lynch *et al*, 2025). Under oxidizing conditions and in absence of DNA, ΔFOSB Cys^172^ and JUND Cys^285^ form a disulfide bond that kinks ΔFOSB (but not JUND), malforming the DNA binding site so that it can no longer bind DNA efficiently (Yin *et al*, 2017; Kumar *et al*, 2022; Lynch *et al*, 2025). ΔFOSB homomers form *in vitro* as well and bind to AP1-consensus DNA sites selectively, but they may not be as sensitive to redox control, and their role *in vivo* is not known (Yin *et al*, 2020). Despite much recent progress, in particular in terms of structure-function relationships, the exact molecular mechanisms of ΔFOSB action remain unclear. Importantly, the exact impact of ΔFOSB on the expression of each target gene, and its broader therapeutic value as a drug target, has yet to be elucidated, despite clear evidence that ΔFOSB serves a critical role in the pathogenesis of several neurological and psychiatric disorders.

Compounds that target ΔFOSB *in vivo*, either by downregulating or upregulating its action, would be enormously useful to interrogate ΔFOSB’s functions and its molecular mechanisms and to elucidate its roles in different disease pathologies and its utility as a therapeutic target. We previously identified JPC0661 as an inhibitor of ΔFOSB action *in vitro* and *in vivo*. JPC0661 is a small molecule that belongs to the 2-phenoxybenzenesulfonic acid-containing class of compounds (McNeme *et al*, 2025). JPC0661 inhibits the binding of the ΔFOSB/JUND heterodimer to a TAMRA-labeled AP1 oligo with IC_50_ ∼11 μM and the ΔFOSB homomer with IC_50_ ∼12 μM (McNeme *et al*, 2025). In cell-based reporter assays, JPC0661 inhibits transcription of a luciferase reporter gene with AP1 consensus sequences built into its promoter region with an IC_50_ less than ∼1 μM (McNeme *et al*, 2025). JPC0661 binds in a groove outside of the DNA binding cleft in the ΔFOSB/JUND bZIP+JPC0661 crystal structure, where it is poised to control DNA-binding and release (**Fig. 1a**) (McNeme *et al*, 2025). Two molecules of JPC0661 are accommodated in a deep hydrophobic groove of the ΔFOSB subunit (Lig1 and Lig2), formed by the side chains of Lys^171^, Arg^175^, Glu^178^, Leu^179^, Arg^182^, and Leu^183^, with the positively charged side chains Lys^171^ and Arg^175^ at one end of the cleft and Arg^182^ at the opposite end (**Fig. 1b**) (McNeme *et al*, 2025). Importantly, the sulfonic acid moiety of JPC0661 Lig1 recruits and rearranges Lys^171^ and Arg^175^, residues that bind to the phosphate-backbone of DNA in the ΔFOSB/JUND+DNA complex and are presumably critical for DNA binding, while the sulfonic acid moiety of JPC0661 Lig2 interacts with ΔFOSB Arg^182^ near the ΔFOSB/JUND interface (**Fig. 1b**) (Yin *et al*, 2017; McNeme *et al*, 2025). JPC0661 is a promising lead compound because it is minimally toxic in Neuro 2A and human neural progenitor cells (McNeme *et al*, 2025). Furthermore, *in vivo* administration of JPC0661 for 3 days into the dorsal hippocampus of APP mice, a mouse model of Alzheimer’s disease neuropathology, reduces the number of ΔFOSB-bound AP1 sites by ∼60% in CUT&RUN studies (McNeme *et al*, 2025). These APP mice express dramatically elevated levels of ΔFOSB protein in the hippocampus and exhibit spontaneous non-convulsive seizure activity similar to that observed in individuals with AD (Corbett *et al*, 2017; You *et al*, 2017). However, despite the encouraging results using JPC0661 as an inhibitor of ΔFOSB action, this compound has several drawbacks. JPC0661 displays only moderate activity *in vivo* (∼60% loss of ΔFOSB DNA-bound sites), two molecules bind side-by-side to the ΔFOSB bZIP subunit in the crystal structure, and the compound uses an induced-fit binding mechanism that rearranges side chains lining the druggable groove. Thus, while JPC0661 demonstrates proof-of-principle that ΔFOSB (and by extension other AP1 transcription factors) can be pharmacologically controlled *in vivo*, it is unknown whether such a lead compound can be further optimized using targeted medicinal chemistry.

**Figure 1.**
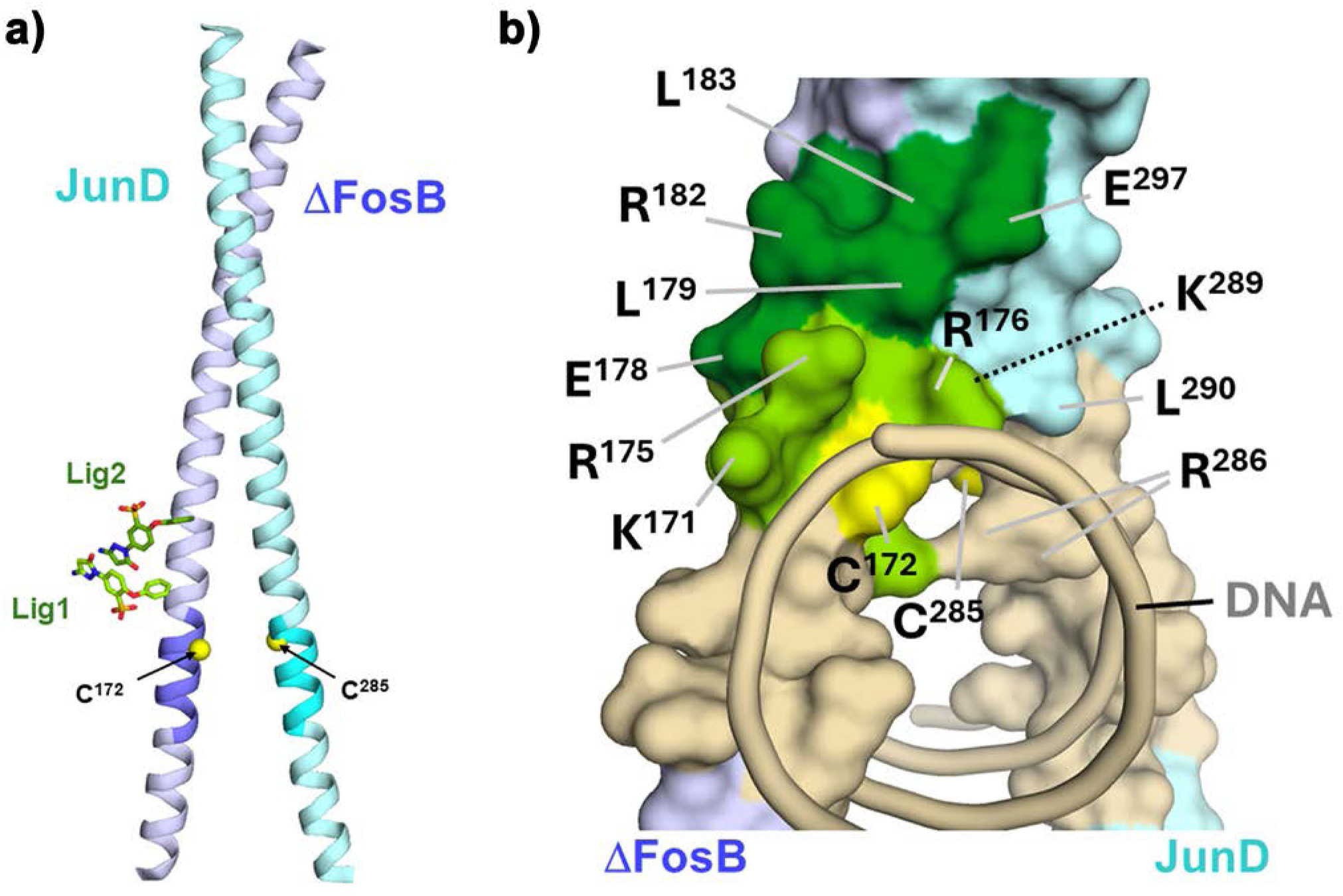
Binding site of the lead compound, JPC0661, on the ΔFOSB/JUND bZIP heterodimer. **A)** 1.7 Å crystal structure of the ΔFOSB/JUND bZIP in complex with JPC0661 (PDB ID: 9OC3). The redox-switch cysteine residues, ΔFOSB C^172^ and JUND C^285^ are highlighted as yellow spheres. ΔFOSB is colored lilac, and JUND is shown in light cyan. Their respective DNA-binding motifs (ΔFOSB N^165^–R^173^ and JUND N^278^–R^286^) are depicted in dark lilac and dark cyan. The JPC0661 ligand molecules (Lig1 and Lig2) are rendered with carbon atoms in light and dark green, respectively; oxygen atoms are in red, nitrogen in blue, and sulfur in yellow. **B)** Overlap of the JPC0661 binding site with residues involved with DNA binding in the ΔFOSB/JUND bZIP mapped onto the ΔFOSB/JUND bZIP+DNA structure (PDB ID: 5VPE). ΔFOSB is shown in lilac and JUND in light cyan. The DNA-binding regions and DNA are shown in tan; the JPC0661-binding region is in dark green, while the overlapping area between the JPC0661-binding and DNA-binding regions is depicted in light green (i.e. residues that contact both JPC0661 and DNA); the redox switch residues, ΔFOSB C^172^ and JUND C^285^, are marked in yellow and also fall within the DNA-binding region. Prominent residues are labeled: ΔFOSB (K^171^, C^172^, R^175^, R^176^, E^178^, L^179^, R^182^, L^183^) and JUND (C^285^, R^286^, K^289^, L^290^, E^297^).

Here, we demonstrate that JPC0661, a first-generation inhibitor of ΔFOSB with cellular and *in vivo* activity, can be optimized using targeted medicinal chemistry, revealing structure-activity relationships that govern compound activity. The resultant analogs display varying abilities to disrupt DNA-binding of ΔFOSB/JUND and ΔFOSB/ΔFOSB biochemically and to work as inhibitors of ΔFOSB-mediated transcription in cell-based luciferase reporter assays. Our best optimized compound, YL0441, shows an IC_50_ < ∼0.1 μM in cell-based reporter assays. Importantly, YL0441 decreases the number of ΔFOSB-bound AP1 peaks by ∼94% in the hippocampus of APP mice *in vivo* and possesses good pharmacological properties, including low cellular toxicity and high metabolic stability. Furthermore, our biochemical studies reveal that YL0441 does not prevent or promote closure of the redox switch, suggesting that its inhibitory effect on DNA-binding, like JPC0661, which binds outside of the DNA-binding cleft, works via a mechanism independent of the redox switch. Our study establishes that 2-phenoxybenzenesulfonic acid-containing compounds can be chemically optimized to yield highly efficacious *in vivo* inhibitors of ΔFOSB function.

## RESULTS

### Strategy to improve JPC0661 as an inhibitor of ΔFOSB/JUND and ΔFOSB/ΔFOSB

Using JPC0661 as a starting lead compound, we developed a medicinal chemistry campaign to optimize JPC0661 for cell-based assays and *in vivo* animal studies (**Fig. 2a** and **Fig. 2b**). JPC0661 is composed of a central phenyl ring with a phenoxy group, a sulfonic acid group, and an amino-pyrazolone group attached. In the JPC0661 co-crystal structure with ΔFOSB/JUND, two molecules bind side-by-side with the phenoxy group of both molecules burying deeply into the hydrophobic cleft of ΔFOSB, anchoring JPC0661 into the compound binding site (McNeme *et al*, 2025). The sulfonic acid group of Lig1 forms a critical salt-bridge and captures DNA-binding residues, ΔFOSB Lys^171^ and Arg^175^, while additional stabilization is provided by π–π stacking and a water-mediated hydrogen-bonding network; the sulfonic acid group of Lig2 interacts with ΔFOSB Arg^182^ (McNeme *et al*, 2025). Guided by a structure-based design, we focused primarily on replacing the amino-pyrazolone moiety with a ring truncation approach (**Fig. 2a**) and alternative substituents such as cinnamamido, benzamido, pyrrolyl, and triazolyl groups, while leaving the phenoxy and sulfonic acid groups anchors intact (**Fig. 2b**). These substitutions were selected based on predicted binding interactions from the ΔFOSB/JUND bZIP+JPC0661 co-crystal structure: amido groups (e.g., cinnamamido, benzamido) were expected to enhance hydrogen bonding at the rim of the cleft, while the heteroaryl caps (e.g., pyrrolyl, triazolyl) could engage in π-π -stacking with nearby aromatic residues. In parallel, a fluoro group was introduced either by replacing the phenoxy group in select compounds (QB0360, QB0361, QB0363, QB0365), which could be leveraged for PET tracer design, or by incorporating fluorinated aryl substituents into the cap region (e.g., 4-F, 4-CF₃) to modulate electronic properties and lipophilicity (QB0301, QB0309) (**Fig. 2b**). The synthetic routes and experimental procedures generating these molecules are provided as Supplemental Information (**Supplemental Material**). The chemical structures and purity of all designed and synthesized new compounds have been validated using a combination of ^1^H NMR, ^13^C NMR, high-resolution mass spectrometry, and HPLC analyses to assure a satisfactory purity prior to biological studies. The resulting panel of 22 analogs of lead compound JPC0661 is also shown in **Supplemental Material Table S1**.

**Figure 2.**
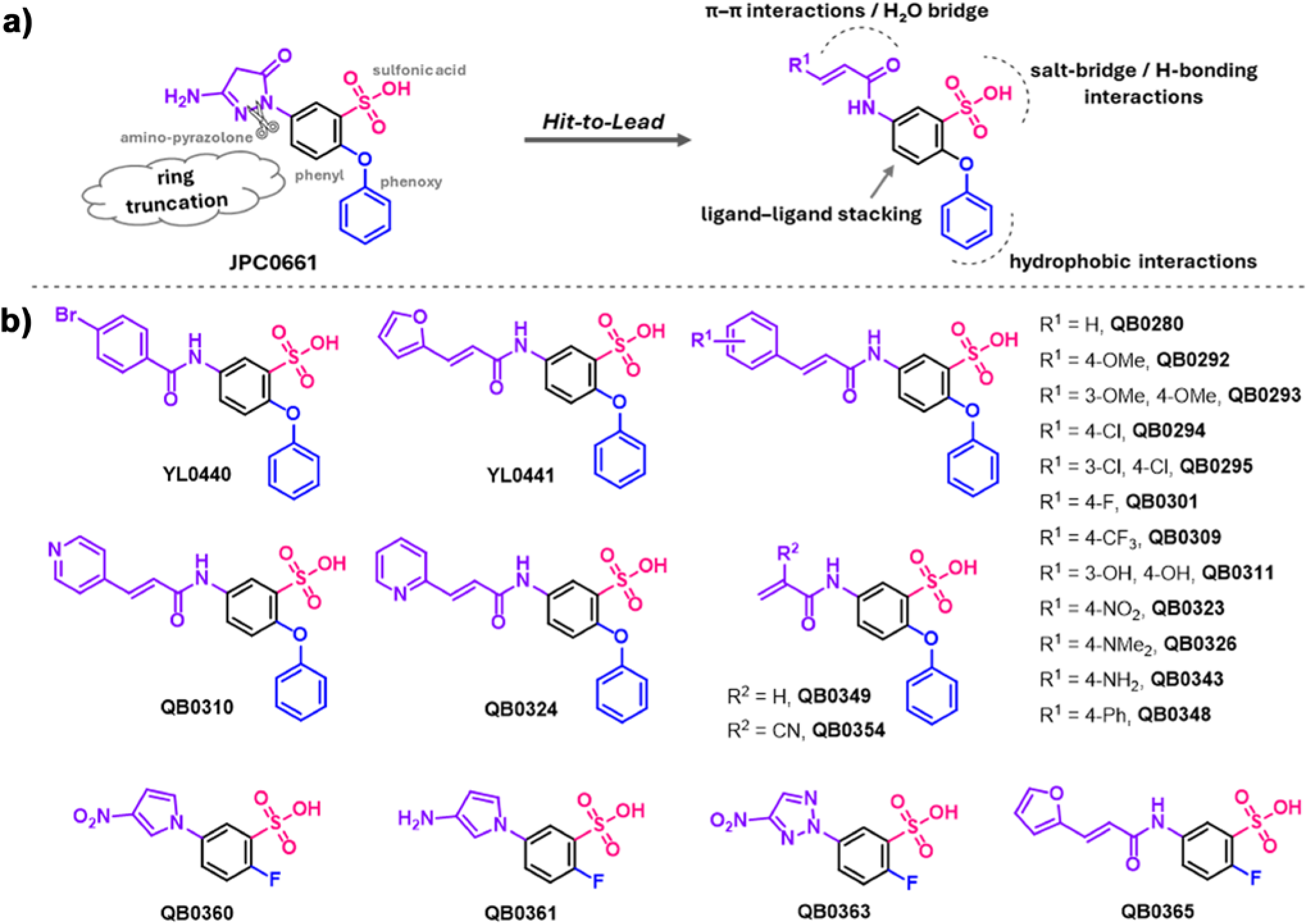
Chemical analogs of JPC0661. **A)** Hit-to-Lead optimization strategy based on the starting lead compound, JPC0661, with chemical moieties named. **B)** Substituents used to probe the role of the phenoxy group and amino-pyrazolone moiety in JPC0661.

### Analogs of JPC0661 disrupt ΔFOSB/JUND and ΔFOSB/ΔFOSB binding to DNA

To assess the activity of our analogs, we used a series of biochemical and cell-based assays. We first measured fluorescence polarization (FP)-based dose response curves (FP-DRC), testing the ability of each compound to disrupt the binding of purified full-length ΔFOSB/JUND heterodimers or ΔFOSB homomers to a TAMRA-labeled oligonucleotide carrying an AP1-consensus site (5’-TGA C/G TCA-3’) (Wang *et al*, 2012; Kumar *et al*, 2022; McNeme *et al*, 2025). The parent compound, JPC0661, inhibited DNA binding to ΔFOSB/JUND and ΔFOSB/ΔFOSB in the micromolar range with IC_50_ 8.9 μM (95% CI 5.7-13.9 μM) and 14.8 μM (95% CI 10.7-20.4 μM), respectively (**Fig. 3a**), as we reported recently (McNeme *et al*, 2025). Out of the 22 compounds, six compounds were considered active with an IC_50_ better than ∼100 μM, namely YL0441, QB0309, QB0348, QB0311, QB0343 and QB0326 (**Fig. 3a**; **Suppl. Fig. S1**). Of these, four compounds looked particularly promising, YL0441 (IC_50_ 13.7 μM and 12.3 μM), QB0309 (IC_50_ 19.1 μM and 30.6 μM), QB0348 (IC_50_ 8.2 μM and 9.4 μM), and QB0311 (IC_50_ 2.9 μM and 10.2 μM) for ΔFOSB/JUND and ΔFOSB/ΔFOSB, respectively (**Fig. 3a**; **Suppl. Fig. S1**). YL0441, QB0309, QB0311, and QB0348 contain an acrylamide linker to the central phenyl ring, attached to furan, 4-(trifluoromethyl)phenyl, 3,4-dihydroxyphenyl, and 4-biphenyl moieties, respectively, instead of the amino-pyrazolone group attached to the central phenyl ring, in addition to the phenoxy and sulfonic acid groups. Twelve compounds had very low activity or were inactive, while four compounds (QB0292, QB0293, QB0294, and QB0295) gave ambiguous results because they altered the fluorescence of the TAMRA-labeled oligonucleotide in absence of purified protein, thus interfering with the assay (**Fig. 3b**; **Suppl. Fig. S1**). Importantly, YL0441, QB309, QB0348, and QB0311 disrupted the DNA-binding of ΔFOSB/JUND heterodimers and ΔFOSB/ΔFOSB homomers similarly well in the FP-assays (**Fig. 3a**) indicating their potential to inhibit various forms of ΔFOSB *in vivo* regardless of its partner, and consistent with JPC0661 binding to ΔFOSB but not JUND in the crystal structure. Thus, four compounds out of the 22-analog panel disrupted DNA-binding biochemically in the low micromolar range.

**Figure 3.**
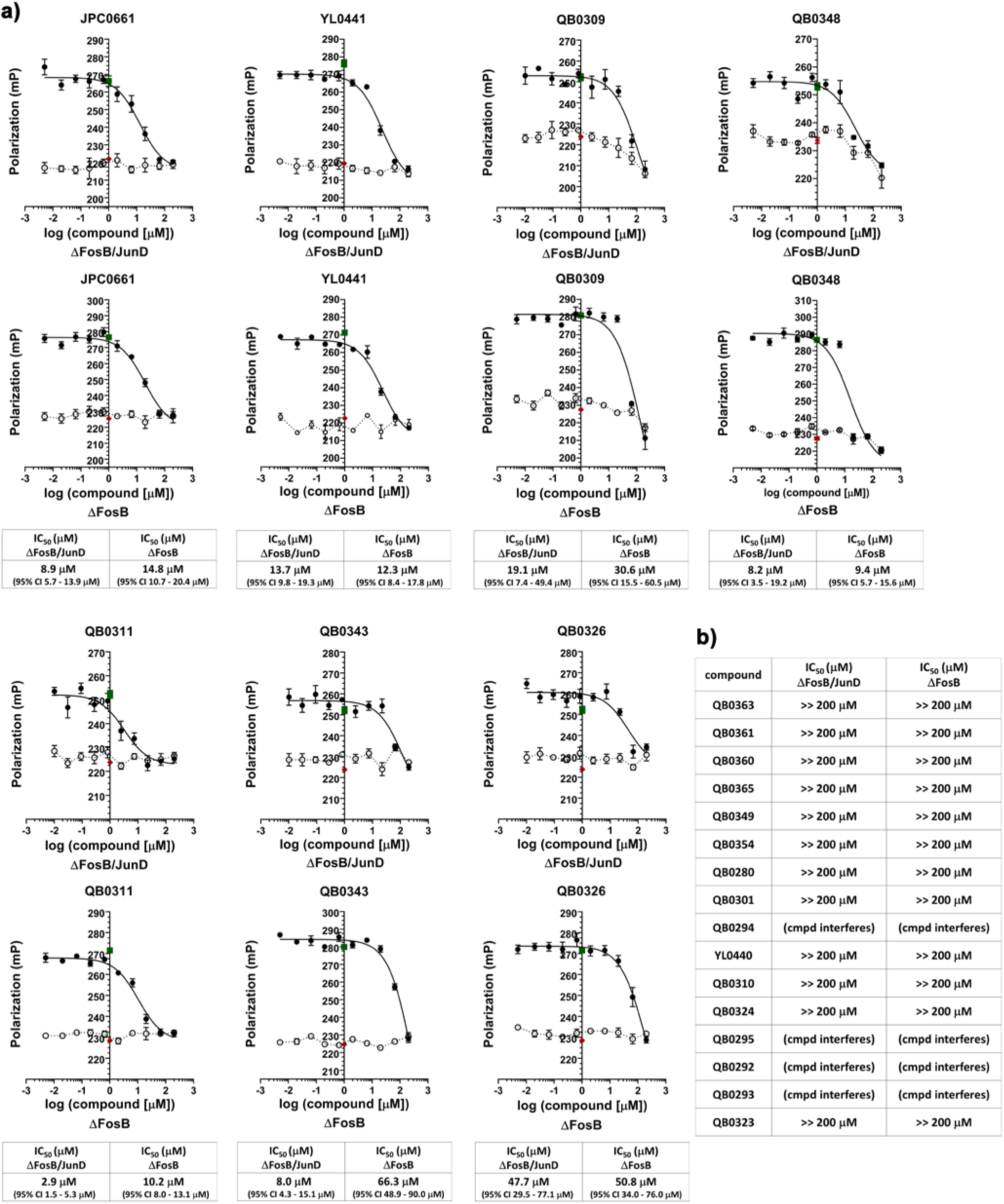
Inhibitory effect of 22 analogs of JPC0661 on DNA binding to ΔFOSB/JUND heterodimers and ΔFOSB/ΔFOSB homomers in fluorescence polarization dose-response curves. **A)** Fluorescence polarization dose-response curves (FP-DRCs) of a TAMRA-labeled oligonucleotide containing the AP1 consensus motif (TMR-cdk5) binding to protein in the presence of varying amounts of compound. In these assays, 25 nM TMR-cdk5 was incubated with either 280 nM ΔFOSB/JUND full-length protein, 320 nM ΔFOSB full-length protein (λ), or no protein (○), along with increasing concentrations of compound (0–200 μM). Control conditions representing 100% inhibition (TMR-cdk5 alone; ♦; n = 16) and 0% inhibition (ΔFOSB/JUND + TMR-cdk5; ▪; n = 16) were also included. The data were fitted with a three-parameter logistic function constraining the bottom of the curve to the TMR-cdk5 oligo alone control (solid lines), yielding estimates for the IC_50_ value that represents the ability of a compound to disrupt protein:DNA-binding. Error bars represent the standard error of the mean (SEM) from four replicate measurements per data point. IC₅₀ values are presented with 95% confidence intervals (CI) for representative plots. **B)** List of the compounds from the 22 analog panel whose effect on DNA binding was limited or which interfered with the assay, interacting directly with the TAMRA-labeled oligonucleotide (“cmpd interferes”), yielding ambiguous results. These compounds were excluded from further investigation.

Next, we tested our panel of compounds for their ability to regulate ΔFOSB-driven gene expression in a cell-based reporter assay (Wang *et al*, 2012; Kumar *et al*, 2022; McNeme *et al*, 2025). We used a luciferase reporter gene under control of AP1-responsive elements stably integrated into HEK293 cells (AP1-luc HEK293 cells) to monitor the effect of compounds on the expression of luciferase (**Fig. 4a**; **Suppl. Fig. S2**). To induce ΔFOSB protein accumulation, these cells were first serum starved (0.5% serum, 24 h) and then stimulated with high serum conditions (20% serum, 24 h), as described previously (Kumar *et al*, 2022; McNeme *et al*, 2025). Our starting compound, JPC0661, inhibited AP1-transcription factor-mediated expression of the luciferase reporter gene with an IC_50_ of ∼0.2 μM in these assays (**Fig. 4a**), somewhat better than we had observed previously (IC_50_ of ∼1-3 μM) (McNeme *et al*, 2025). Of the six compounds most active in the biochemical FP-DRC assay (which we set as a necessary prerequisite to consider a compound), three compounds were also robustly active in the reporter assay, namely, YL0441 (IC_50_ 0.10 μM, 95% CI 0.01-1.85 μM), QB0309 (IC_50_ 0.64 μM, 95% CI 0.14 - 2.84 μM) and QB0348 (IC_50_ 0.63 μM, 95% CI 0.05 - 7.55 μM) (**Fig. 4a**). Compounds QB0311, QB0343, and QB0326 were minimally inhibiting or not at all (**Fig. 4a**). Several compounds showed significant inhibitory effect on gene expression with IC_50_ values less than ∼1 μM in our reporter assay, despite being minimally active or inconclusive in the biochemical FP-DRC assay (e.g., QB0349, QB0354, QB0301, QB0294) (**Suppl. Fig. S2**). This highlighted the experimental gap between testing compounds with isolated, purified AP1 transcription factors in aqueous solution and testing them in a cellular context where ΔFOSB protein (and any other endogenous AP1 transcription factors) are induced via serum stimulation when the complete endogenous transcription machinery is present. To assess the toxicity of the compounds under conditions similar to the reporter assay, we tested cell viability as a function of increasing compound concentration (0-100 μM) with the CellTiter-Glo viability assay in AP1-luc HEK293 cells (**Fig. 4b**; **Suppl. Fig. S3**). Of the compounds that were the most active as inhibitors of ΔFOSB function both in our biochemical assay as well as in our cellular assay, i.e., YL0441, QB0309, and QB0348, only YL0441 showed no noticeable decrease in cell viability (**Fig. 4b**). Overall, out of the 22 analogs tested, 11 showed no overt decrease in cell viability (at 31.5 μM), indicating that our series of 2-phenoxybenzenesulfonic acid analogs appears generally well tolerated in cells, serving either as active compounds or inactive control compounds for future *in vivo* studies (**Fig. 4b**; **Suppl. Fig. S3**).

**Figure 4.**
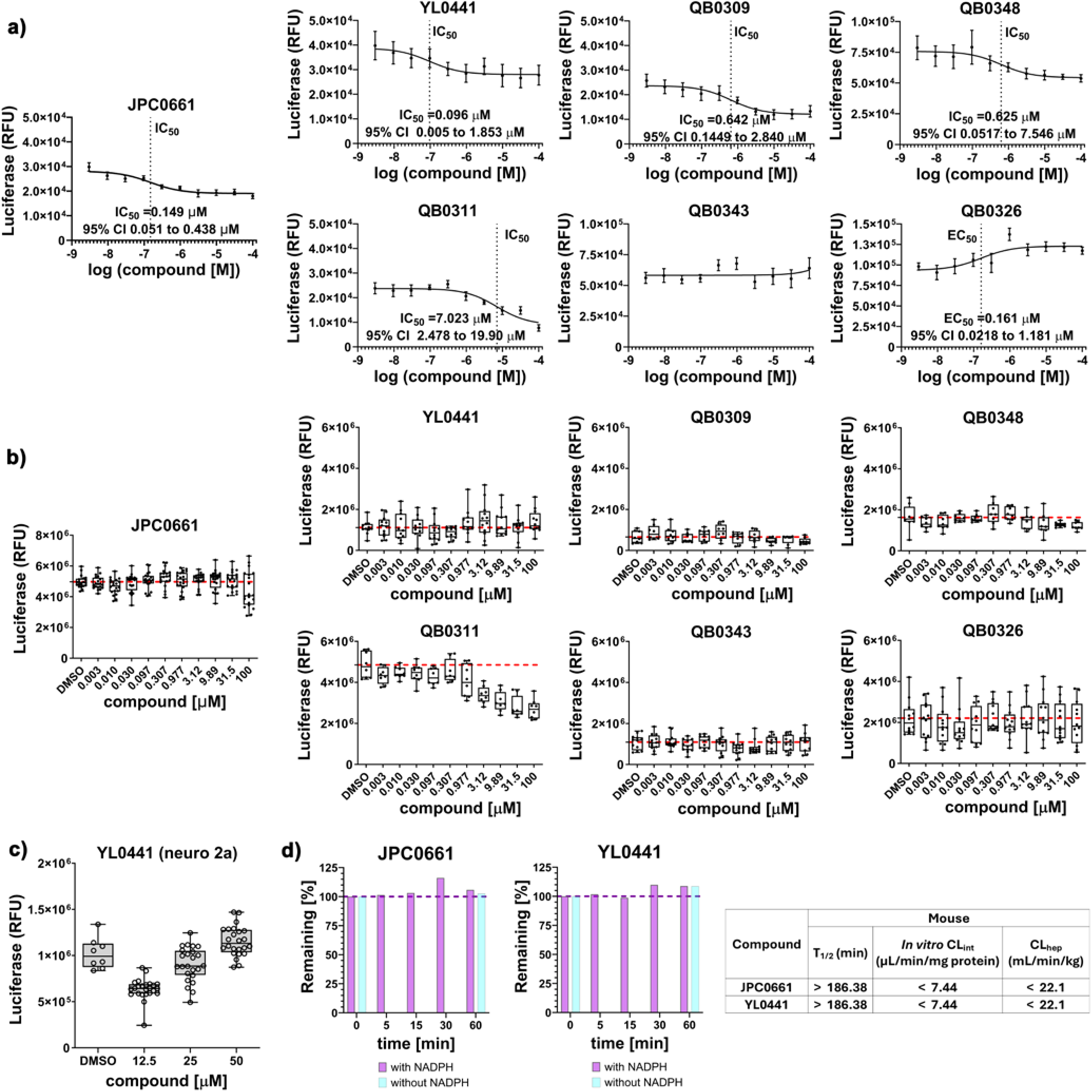
Analogs of JPC0661 regulate expression of an AP1-driven reporter gene in cell-based assays. **A)** Effects of compounds (0–100 μM) on AP1-luciferase reporter activity evaluated in AP1-luc HEK293 cells. Dose-dependent activation of the AP1 reporter was quantified by measuring changes in luciferase signal, expressed as relative fluorescence units (RFU). Each compound was tested in at least two independent experiments (n = 4 wells per experiment nominally), combined resulting in at least a total of 6–8 wells per compound concentration, and the results normalized to the luciferase signal from blank wells within each experiment (n = 8 wells). Data were fitted to a three-parameter logistic model to determine IC_50_ values and associated 95% confidence intervals (CI). Data points are presented as mean ± SEM. **B)** Effect of serially diluted compounds (0.003–100 μM) on the viability of AP1-luc HEK293 cells assessed using the CellTiter-Glo viability assay (2-hour incubation). Cell viability was normalized to the control containing 0.5% (v/v) DMSO in the absence of compound (n = 7 wells). Data points represent the mean ± SEM, with a total of 10–12 wells analyzed per concentration. A red dotted line indicates the average of the data points for 0.5% (v/v) DMSO alone, no compound. **C)** Effect of YL0441 on the viability of Neuro2a cells assessed using the CellTiter-Glo cell viability assay after 72-hour incubation at 0, 12.5, 25, and 50 μM compound concentration. Viability was normalized to a DMSO control that contained 0.5% DMSO and no compound (n = 8 wells). Data are shown as mean ± SEM, with 12 wells analyzed per concentration. **D)** The stability of YL0441 versus JPC0661 in mouse liver microsomes was assessed by analyzing compound depletion over time with and without NADPH with LC/MS/MS.

Given its favorable chemical properties as well as promising biochemical and cell-based activity, YL0441 was selected for further studies. To assess the toxicity of YL0441 prior to *in vivo* administration studies, we tested the impact of YL0441 on the viability of Neuro 2A cells over 3 days up to a maximum dose of 50 μM, revealing that YL0441 is well tolerated in these cells (**Fig. 4c**). To assess the metabolic stability of YL0441, we used a mouse liver microsomal stability assay and revealed that YL0441 is at least as stable as our starting compound, JPC0661, with no apparent loss of compound within the 60-minute timeframe tested (**Fig. 4d**). Thus, leveraging a panel of 22 novel analogs, we successfully generated a new lead compound, YL0441, which inhibits AP1-driven gene expression effectively in a cell-based reporter assay and possesses low toxicity and favorable metabolic stability (summarized in **Supplemental Material Table S1**).

### YL0441 efficiently disrupts ΔFOSB binding to genomic DNA in brain *in vivo*

To test whether YL0441 disrupts the binding of endogenous ΔFOSB to genomic DNA *in vivo*, we infused YL0441 (or vehicle) directly into the hippocampus of APP mice and then assessed the amount of ΔFOSB bound to DNA in the presence or absence of compound. YL0441 or vehicle was loaded into osmotic minipumps and infused for three days via a cannula directly into the right dorsal hippocampus of 2.5-month-old APP transgenic mice. At this age, hippocampal ΔFOSB levels are robustly and consistently elevated in these mice (Corbett *et al*, 2017). CUT&RUN-sequencing was then performed on the dorsal two-thirds of hippocampal tissue receiving infusion. The contralateral (left) hemibrain, which did not receive any infusion, was fixed and immunostained for ΔFOSB using an anti-ΔFOSB antibody (Cell Signaling, #D3S8R) to confirm elevated levels of ΔFOSB in the hippocampus of these APP mice (**Suppl. Fig. S4**).

ΔFOSB-bound chromatin was profiled using an anti-ΔFOSB antibody (Cell Signaling, #D3S8R), and the number of ΔFOSB-bound DNA peaks quantified as described previously (Yeh *et al*, 2023; McNeme *et al*, 2025). Importantly, this anti-ΔFOSB antibody does not bind significantly to brain slices of *FosB/ΔFosB* knockout mice (Yutsudo *et al*, 2013), so that the peaks pulled down represent ΔFOSB bound to genomic DNA, and not any other AP1 transcription factors or off-target proteins bound to DNA (Yeh *et al*, 2023). This enabled us to examine the *in vivo* impact of YL0441, directly focusing on ΔFOSB alone. Administration of YL0441 resulted in the dramatic loss in the total number of ΔFOSB binding peaks. Whereas animals treated with vehicle displayed 9,642 peaks, YL0441-treated animals retained only 540 peaks, a 94% reduction in total binding events (**Fig. 5a**). Furthermore, the average peak width decreased from ∼286 bp (vehicle-treated mice) to ∼254 bp (YL0441-treated mice), suggesting a widespread loss of ΔFOSB-chromatin engagement and reduced ΔFOSB binding (**Fig. 5b**). This findings indicate that YL0441 interferes with ΔFOSB binding to DNA and/or promotes its release from chromatin in vivo.

**Figure 5.**
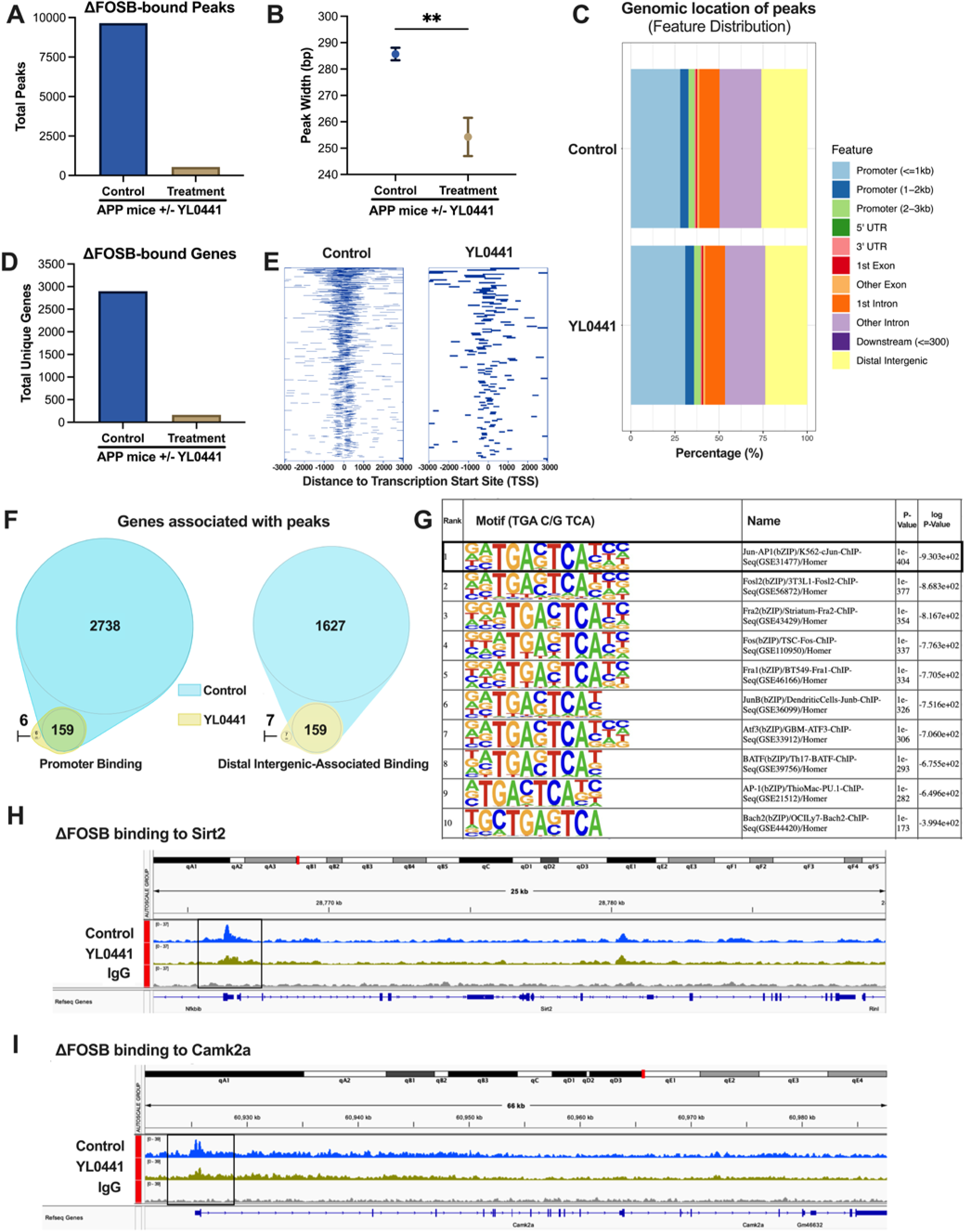
YL0441 potently disrupts ΔFOSB–chromatin engagement in the dorsal hippocampus of APP mice. YL0441 (‘Treatment’) or vehicle (‘Control’) was infused via a cannula unilaterally into the dorsal hippocampus of APP mice for 3 days using osmotic minipumps. CUT&RUN-sequencing was performed on nuclei isolated from the dorsal hippocampal tissue of each individual animal treated with either vehicle or YL0441 to determine the genome-wide binding of ΔFOSB to genomic DNA *in vivo*. **A)** Total number of high-confidence ΔFOSB-bound peaks detected in vehicle-infused (‘Control’) versus YL0441-infused (‘Treatment’) mice. **B)** Average peak width (bp) of the ΔFOSB-bound peak sets in **A)**. Data for **A)** and **B)** below are shown as mean ± SEM; **p < 0.01. No sex differences were observed for **A)** and **B)**. **C)** Genomic distribution of the ΔFOSB-bound peaks under vehicle and YL0441 conditions reveals a widespread distribution over promoter, intronic, and intergenic regions in the dorsal hippocampus, with similar patterns seen in Treatment and Control conditions. Stacked bar charts depict the different classes of genomic features occupied by ΔFOSB (UCSC annotations), including the regions promoter-proximal (≤ 1 kb) and promoter-distal (1–5 kb) with respect to the translation start site (TSS), the 5′ UTR/first exon, intragenic (internal exons + introns) regions, the 3′ UTR/downstream region, and distal intergenic regions. Values are expressed as percentages of the total peaks in each condition (Control vs YL0441-treatment). **D)** Total number of unique ΔFOSB-bound genes in vehicle-infused vs. YL0441-infused APP mice shows an approximately 94% reduction in ΔFOSB binding after YL0441 treatment. **E)** ChIPseeker heatmaps showing the peak distribution of ΔFOSB binding intensities centered around the TSSs in APP mice under vehicle-(left) and YL0441-(right) infused conditions. YL0441 results in a dramatic loss of ΔFOSB binding near TSS sites at gene promoters. Note: each bar represents one ΔFOSB CUT&RUN peak plotted relative to its genomic distance to the nearest annotated TSS along the X-axis. The bar’s length shows the span of that peak (in bp). Negative values indicate upstream locations from the TSS (i.e., ‘promoter’), positive values indicate downstream locations (i.e., gene body/5′ UTR), with the TSS defined at x=0. Peaks are stacked from top-to-bottom vertically, sorted by the distance to the TSS, so that the top of the stack lists genes with the most ΔFOSB-peaks bound within −3000 to +3000 bp from the TSS, and the bottom, the least. **F)** Number of unique genes with ΔFOSB peaks bound to promoter regions, i.e., falling within −3000 to +3000 bp of the TSS (left) compared to those bound to distal intergenic regions (right). Venn diagram analysis reveals that the predominant effect of YL0441 (gold) is to globally deplete peaks seen under vehicle-treated conditions (light blue), regardless of their location within the chromatin. **G)** De-novo motif analysis showing that the vast majority of ΔFOSB-bound peaks (in vehicle-infused animals) are bound to DNA sequences that contain the AP1 consensus (TGA C/G TCA). **H)** Example of ΔFOSB binding at the promoter region of *Sirt2* promoter (Genome browser snapshot of chr7: position) illustrating the dramatic loss of ΔFOSB occupancy upon YL0441 exposure. The tracks are shown for samples from the dorsal hippocampus of APP mice infused with vehicle (top track in sky blue) or with YL0441 (middle track in gold-green) using an anti-ΔFOSB-antibody for CUT&RUN-sequencing vs. using an IgG-antibody as a control for non-specific binding (bottom track in grey; subtracted background). **I)** Example of ΔFOSB binding at a distal intergenic/putative enhancer region of an upstream intergenic enhancer (12.5 kb) at the *Camk2a* locus (Genome browser snapshot of chr18: position), demonstrating a dramatic loss of ΔFOSB-bound sites in response to YL0441-treatment. Track conventions as described in **H)**.

Genomic annotation revealed that, in vehicle-treated mice, ΔFOSB peaks were broadly distributed across promoters (34%), exons (10%), introns (35%), and distal intergenic regions (21%) (**Fig. 5c**). After treatment with YL0441, the collection of residual peaks displayed a similar distribution (**Fig. 5c**). The ΔFOSB-binding peaks at promoters of target genes (i.e., located <300 kb from a transcription start site (TSS)) mapped to ∼ 2,897 unique genes in vehicle-treated mice, which decreased to ∼165 genes in YL0441-treated mice (**Fig. 5d**). ChIPseeker heatmaps showed that under normal conditions (i.e., vehicle-treated mice) ΔFOSB-peaks clustered tightly in a vertical column centered on 0 bp (i.e., the TSS of genes), indicating ΔFOSB binding at promoters. Many fewer binding sites were observed at ±1–3 kb distance from the TSSs (either 5’-upstream or 3’-downstream), indicating a much smaller number of ΔFOSB peaks that bound to such proximal regulatory sites (**Fig. 5e**). More distal regulatory peaks, located far from TSSs, were also lost, indicating a universal decrease in ΔFOSB-binding sites on a broad scale as a result of YL0441-treatment. The remaining few ΔFOSB peaks clustered predominantly within ∼500 bp of TSSs, suggesting perhaps a small cohort of high-affinity sites that YL0441 fails to dislodge completely (**Fig. 5e**). Depicting these results in Venn diagrams shows that, of the 2,897 unique genes associated with promoter peaks (i.e., 2738+159), only 165 genes were left after treatment with YL0441 (**Fig. 5f**). Likewise, of the 1,786 unique genes associated with distal intergenic peaks (i.e., 1627+159), only 166 genes were left after treatment with YL0441 (**Fig. 5f**). Importantly, we found no evidence that YL0441 treatment induces ΔFOSB binding at sites not occupied under control conditions.

*De novo* motif analysis confirmed that the large majority of ΔFOSB peaks from vehicle-treated mice (9642 ΔFOSB peaks) were enriched for the canonical AP-1 binding motif (TGAC/GTCA), consistent with specific ΔFOSB occupancy (**Fig. 5g**). Analysis of single representative loci from canonical ΔFOSB targets revealed large-scale YL0441-induced depletion of ΔFOSB peaks, for instance, at the *Sirt2* and *Camk2a* loci – two known in vivo targets of ΔFOSB (Teague & Nestler, 2022) – in IGV Browser snapshots (**Fig. 5h** and **5i**). Collectively, these data indicate that YL0441 dramatically reduces the abundance and intensity of ΔFOSB–DNA interactions *in vivo*, erasing 94% of chromatin contacts within the 72-hour duration of compound administration.

### Insight into the mechanism of YL0441 action

To gain insight into the molecular mechanism of YL0441 action, we probed its effect on the redox switch within ΔFOSB/JUND heterodimers: ΔFOSB Cys^172^/JUND Cys^285^. Closure of the redox switch (e.g., in response to oxidative stress) on the one hand induces a dramatic conformational change of ΔFOSB (kinking the helix in two at the hinge Arg^176^-Arg^177^) so that ΔFOSB Cys^172^ can approach and form a disulfide bond with JUND Cys^285^. On the other hand, it dramatically inhibits DNA binding by malforming the DNA-binding cleft (Yin *et al*, 2017; Kumar *et al*, 2022; Lynch *et al*, 2025). To assess whether binding of YL0441 might alter the ability of the bZIP helices to splay open and insert their DNA-binding motifs into the major groove of DNA, we tested whether the addition of YL0441 alters the flexibility of the ΔFOSB bZIP alpha-helix in general, using closure of the redox switch as a read-out. To do this, we oxidized recombinant ΔFOSB/JUND bZIP (containing the sole cysteine residues ΔFOSB Cys^172^ and JUND Cys^285^) with diamide (100 μM) in the presence of excess compound (500 μM) and monitored whether the redox switch was able to still close, generating a disulfide-bonded heterodimer. We compared YL0441, JPC0661 (our starting lead), and the inactive compound QB0365 in this biochemical assay. As a control, the ΔFOSB/JUND bZIP was treated with diamide alone (no compound) to close the redox switch. We also treated the protein with Z21599131480, which covalently modifies ΔFOSB Cys^172^, rendering the redox switch constitutively open (Kumar *et al*, 2022) as a second control (not shown). All three compounds (YL0441, JPC0661 and QB0365) did not prevent closure of the redox switch in ΔFOSB, as assessed with semi-native SDS-PAGE (i.e., no reducing agent in the loading buffer) (**Fig. 6a**). We next tested whether these compounds instead could promote oxidation of ΔFOSB/JUND bZIP directly, inducing disruption of DNA-binding in this way, and revealed that YL0441, JPC0661, and QB0365 did not oxidize the protein and promote closure of the redox switch (**Fig. 6b**). Thus, the mechanism by which YL0441 disrupts ΔFOSB binding to DNA does not involve the redox switch, consistent with the location of the JPC0661 binding site in a druggable groove outside of the DNA-binding cleft.

**Figure 6:**
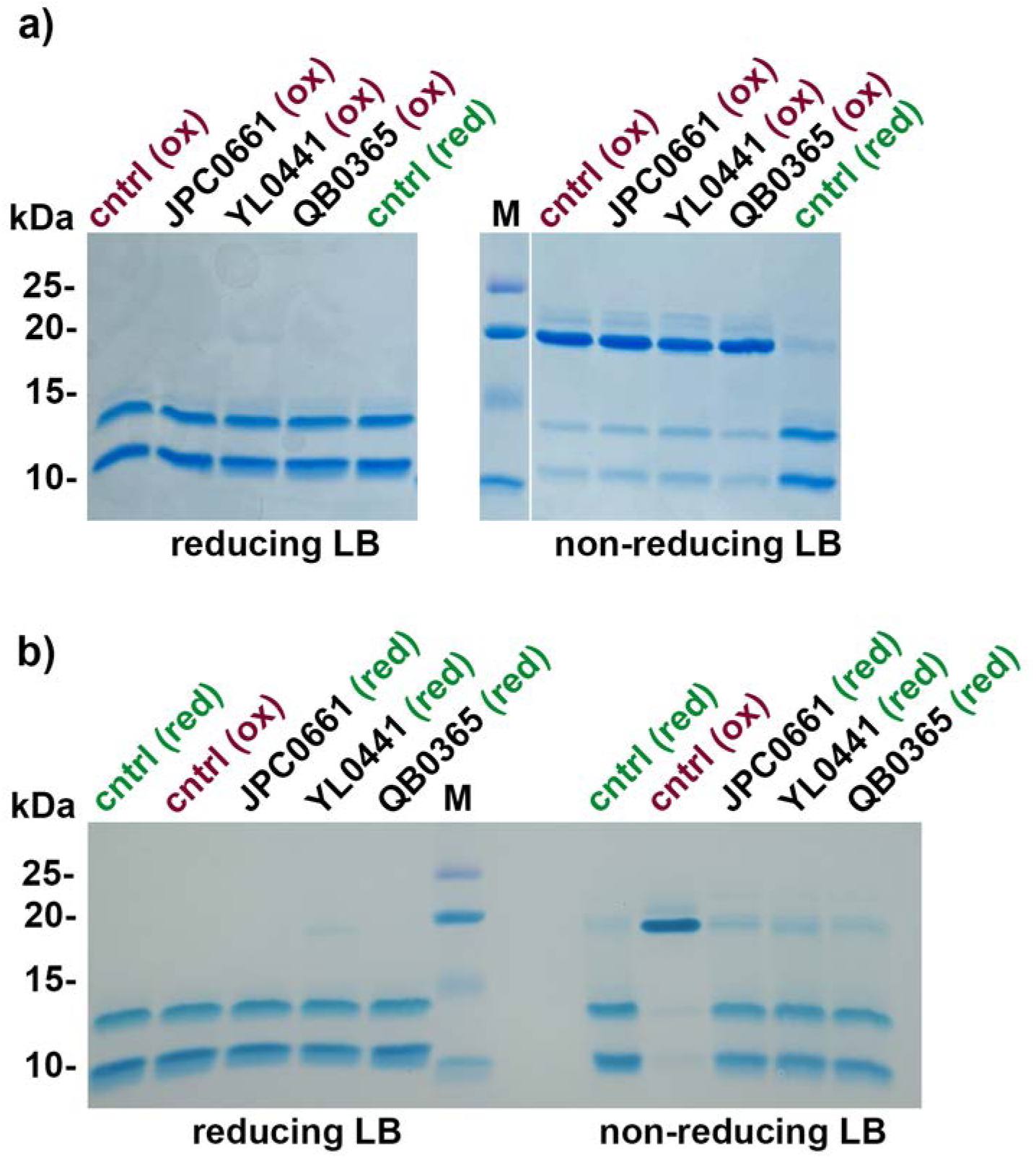
Molecular insight into YL0441 action. **A)** JPC0661, YL0441, and QB0365 do not prevent closure of the redox switch. ΔFOSB/JUND bZIP was incubated with compounds (0.5 mM) with 100 μM diamide (‘ox’, oxidized) or without diamide (‘red’, reduced) and assessed by SDS-PAGE (with or without reducing agent in the loading buffer). **B)** JPC0661, YL0441, and QB0365 do not promote closure of the redox switch by oxidizing the protein. ΔFOSB/JUND bZIP protein was incubated with compounds (0.5 mM) with no diamide (‘red) while protein alone (no compound) was also incubated with diamide (‘ox’; 100 μM) as a control. Samples were then assessed by SDS-PAGE (with or without reducing agent in the loading buffer). In **A)** and **B)** ‘cntrl’ denotes the ΔFOSB/JUND bZIP protein in absence of compound; ‘M’ indicates the molecular markers (kDa).

## DISCUSSION

Here, we demonstrate, starting from a lead, JPC0661, that we can chemically optimize 2-phenoxybenzenesulfonic acid-containing compounds as inhibitors of ΔFOSB function *in vitro* and *in vivo*. Our best analog, YL0441, has an IC_50_ 0.1 μM in AP1-reporter-assays and leads to an almost complete loss of ΔFOSB-binding sites to genomic DNA in brain *in vivo.* This *in vivo* activity represents a dramatic improvement compared to JPC0661, which triggered only a ∼60% loss of such ΔFOSB-bound sites (McNeme *et al*, 2025). YL0441 is metabolically stable and displays no detectable cellular toxicity, suggesting that it is a promising starting point for further development and can be leveraged to assess the utility of ΔFOSB as a therapeutic target in animal models for an array of different neuropsychiatric disorders. Furthermore, biochemical and cellular screening of our panel of analogs permits us to establish a preliminary structure-activity relationship (SAR) so that we can design further optimized chemical probes and diagnostic tools suitable for an array of different *in vivo* studies and uses.

### Novel compounds targeting ΔFOSB transcription factor action

Despite their clear involvement in many human diseases, relatively modest progress has been made in targeting the AP1 transcription factor family pharmacologically (Ye *et al*, 2014; Brennan *et al*, 2020; Casalino *et al*, 2022). This family is composed of many members, in addition to ΔFOSB and FOSB, including FOS, FOSL1, FOSL2, JUN, JUNB, and JUND, as well as related bZIP transcription factors such as the ATF family members and CREB family members (Bejjani *et al*, 2019). These proteins form heterodimers, and in some cases homomers, via their DNA-binding bZIP domains in a mix-and-match strategy (Lambourne *et al*, 2025). Importantly, the different AP1 transcription factor complexes each have unique biological functions (Bejjani *et al*, 2019; Song *et al*, 2023). Thus, compounds that discriminate between the different AP1 subunits and bind selectively to a particular subunit would be tremendously valuable. However, the DNA binding cleft buried deep in the fulcrum of bZIP dimer forceps, an obvious site for drug discovery, is very highly conserved among the different AP1-subunits (Kumar *et al*, 2022). We have shown that the redox switch residue, ΔFOSB Cys^172^, another obvious site for drug discovery, indeed can be covalently modified with small molecules, altering ΔFOSB-driven gene expression in cell-based assays (Kumar *et al*, 2022). However, compounds covalently modifying cysteine residues, though gaining acceptance as a drug discovery strategy and FDA approval, are not easy to develop because of their chemically reactive nature, and greater potential to display off-target effects, metabolic instability, and/or toxicity (Bauer, 2015; Gehringer & Laufer, 2019). Our discovery of a druggable groove within the ΔFOSB subunit, outside of the DNA-binding cleft and with little sequence conservation among AP1 transcription factor family members, offers a radically new strategy to drug this family (McNeme *et al*, 2025). Starting from JPC0661, which binds to this groove, we designed and tested a panel of 2-phenoxybenzenesulfonic acid-containing analogs in this study to further explore and validate this lead compound.

Our medicinal chemistry effort produced the improved inhibitor YL0441, which potently inhibits ΔFOSB function *in vivo* and establishes preliminary structure-activity relationships. The sulfonic acid moiety is indispensable for activity, and it appears to engage DNA-contacting residues ΔFOSB Lys^171^ and Arg^175^ as well as ΔFOSB Arg^182^ in the crystal structure of ΔFOSB/JUND bZIP+JPC0661 at either side of the groove (McNeme *et al*, 2025). The phenoxy group anchors the scaffold deep in the hydrophobic cleft, establishing the core pharmacophore. Therefore, our campaign focused on replacing the amino-pyrazolone cap of JPC0661, because it confers only modest efficacy, whereas its replacement with cinnamamido or heteroaryl substituents (such as biphenyl or furan) produces a marked gain in potency and cellular activity, leading to an excellent inhibitory profile for YL0441. It is important to note that while the biochemical IC_50_ values of YL0441 and our starting compound JPC0661 are very similar in FP-DRCs (∼10-15 μM for ΔFOSB/JUND and ΔFOSB), YL0441 is ∼50-100-fold more active in the cell-based reporter assay (< 0.1 μM) and leads to a near complete loss of ΔFOSB-binding peaks bound to genomic DNA *in vivo* compared to only 60% for the lead molecule. This highlights the importance of leveraging multiple orthogonal assays to home in on compounds that will have the most *in vivo* efficacy. The biochemical FP-assay, though very useful in the early stages of chemical probe development, utilizes recombinant proteins that likely do not fully capture the properties of ΔFOSB as it presents itself in a cellular environment, nor the additional effects and complexities introduced by the presence of binding partners such as co-factors and AP1 consensus sites presented in the context of chromatin, not a short oligonucleotide. In addition to YL0441, QB0309, and QB0348 also exhibited strong activity, though with moderate cellular toxicity, identifying them as possible secondary leads. QB0311 showed weak cellular efficacy and significant cellular toxicity despite its potency in biochemical assays, indicating pharmacokinetic (ADMET) limitations. Notably, several fluorinated analogs (e.g., QB0360 and QB0361), though not measurably active, also did not decrease cell viability, so that these compounds might hold translational interest for their potential development as PET ligands for *in vivo* imaging if their activity can be improved. Thus, the 2-phenoxybenzenesulfonic acid family of compounds forms a chemical scaffold amenable to optimization through rational medicinal chemistry modifications.

### ΔFOSB and other AP1 transcription factors as drug targets

The ability of YL0441 to pharmacologically reduce the occupancy of ΔFOSB to genomic DNA so effectively *in vivo* is striking. Three-day infusion of YL0441 reduced ΔFOSB-binding sites from 9,642 peaks (vehicle) to 540 peaks (compound-treated) via a global release of transcription factor from all AP1-motifs (i.e., at promoter regions, gene bodies, and distal intergenic regions) and not selective pruning of individual loci. While the number of 9,642 ΔFOSB-binding peaks experimentally determined in vehicle-treated tissue may seem large, it likely represents only a fraction of the possible AP-1 motif landscape. In the 3.2-Gb human genome, AP-1 motifs are predicted to occur roughly once every 1.24 × 10⁴ bp, yielding ∼260,000 potential binding sites (Zhou *et al*, 2005). In practice, however, only a small subset of these motifs is experimentally identifiable at a time. On the one hand, inaccessibility due to chromatin, as well as the competition and/or cooperativity of (different AP1) transcription factors can all gate access to these motifs in a cell-type specific and cellular state-dependent manner (Bejjani *et al*, 2019, 2021). On the other hand, ChIP-seq/CUT&RUN experiments inevitably under-sample very transient and/or low-occupancy protein:DNA interactions. Similarly to the studies presented here, a recent study detected 8,199 ΔFOSB AP1 peaks in nucleus accumbens of saline-treated mice, rising to 11,843 ΔFOSB AP1 peaks after chronic cocaine exposure (a ∼44 % increase) (Yeh *et al*, 2023). Thus, our studies using dorsal hippocampal-derived tissue from APP mice treated with vehicle or YL0441 fall well within the expected experimental range. The near-total loss of ΔFOSB bound to genomic DNA *in vivo*, together with the significant narrowing of the residual peak widths, indicates that YL0441 potently reduces ΔFOSB chromatin occupancy and does not permit ΔFOSB to be redirected to ectopic, non-related AP1 motifs. Thus, while these data do not distinguish whether the compound prevents initial DNA engagement or accelerates dissociation of pre-bound complexes, the preservation of canonical AP-1 motif enrichment among the few remaining peaks suggest that the compound promotes a dramatic loss of occupancy and not retargeting to other sites in genomic DNA.

Our studies also highlight the utility of YL0441 to probe the role and mechanism of ΔFOSB as an epigenetic factor *in vivo*. AP1 transcription factors are thought to play important roles working as context-dependent pioneer transcription factors, and proteins that remodel and maintain chromatin and its accessibility to prepare target genes for transcription (Biddie *et al*, 2011; Bejjani *et al*, 2021). In eukaryotes, transcription factors are thought to assemble into large protein complexes at enhancer and promoter sites, that contain multiple transcription factors and other components of the transcription machinery (Veitia, 2025). ΔFOSB likely incorporates into such multiprotein complexes as well, but whether these are all tightly bound to genomic DNA or more loosely is not known. Recent work suggests that AP1 transcription factors take turns occupying the same AP1 sites in genomic DNA called ‘AP1 hotspots’ that are located predominantly at distal intergenic regions (Seo *et al*, 2021; Bejjani *et al*, 2021). The pattern of AP1 hotspots open to recruiting AP1 transcription factors depends on the exact cellular circumstances, so that in response to cell type–specific and cell-environment cues, AP1 transcription factors occupy genomic loci in distinct non-overlapping patterns (Seo *et al*, 2021; Bejjani *et al*, 2021). At these genomic sites, AP1 transcription factor subunits also recruit many different bZIP-containing partners, and work collaboratively to generate a portfolio of various, non-redundant DNA-binding transcription factor complexes which can bind to both shared (overlapping), as well as unique, genomic loci (Xie *et al*, 2025; Fonseca *et al*, 2019; Bejjani *et al*, 2021). Therefore, it is even more impactful that YL0441 is able to disrupt virtually all ΔFOSB-bound AP1-sites. This is an important finding because ΔFOSB, like other AP1 transcription factors, can work either as an activator or repressor of gene expression depending on the exact target gene and cellular context, likely because it can recruit different bZIP dimerization partners or different co-factor proteins (Xie *et al*, 2025; Fonseca *et al*, 2019; Bejjani *et al*, 2021). Because JPC0661 appears to engage only ΔFOSB, not JUND, in the crystal structure, it appears that such a compound or its optimized analogs, could be developed to target a specific AP1 subunit (e.g., ΔFOSB) selectively over other AP1 transcription factors and do so with high efficiency. Such AP1-subunit-specific compounds would be powerful tools to experimentally dissect out the effect of inhibiting a particular AP1 transcription factor *in vivo* (regardless of its particular dimerization partner), for instance, to probe the expression of particular gene targets such as those mediating long-lasting maladaptations underlying drug addiction, cognitive decline, and dyskinesias.

It is of paramount importance to identify and validate a set of compounds which disrupt the binding of ΔFOSB to AP1 sites *in vivo* pharmacologically so that key disease-driving genes can be identified and their relationship to the different disease pathologies delineated. Genetic studies demonstrate that ΔFOSB has unique and controlling effects on the pathogenesis of several neuropsychiatric disorders. Genetically inhibiting elevated ΔFOSB in the striatum in response to drugs of abuse reduces their rewarding effects (Teague & Nestler, 2022; Robison & Nestler, 2022), while decreasing ΔFOSB in response to chronic L-DOPA used to treat Parkinson’s disease patients reduces abnormal involuntary movements (AIMs) while leaving the antiparkinsonian action of L-DOPA undiminished (Beck *et al*, 2021). Genetic inhibition of accumulated ΔFOSB in hippocampus in mouse models of Alzheimer’s disease reverses cognitive deficits in AD mice (You *et al*, 2017; Corbett *et al*, 2017). YL0441 provides a clean chemogenetic tool to probe ΔFOSB-dependent transcriptional programs *in vivo* and to dissect how persistent AP1 signaling shapes gene networks that converge on disease-relevant phenotypes. YL0441 also sets a quantitative benchmark for future ΔFOSB antagonists and raises the possibility of leveraging AP-1 displacement therapeutically in disorders where pathological ΔFOSB accumulation is a driver of maladaptive transcriptional memory.

## ABBREVIATIONS

a.a.: amino acid
Aβ: Amyloid β protein
AD: Alzheimer’s disease
AP1: Activator protein 1
APP: amyloid precursor protein
BSA: bovine serum albumin
bZIP: basic leucine zipper
CI: confidence interval
DMSO: dimethyl sulfoxide
DRC: dose-response curve
DMEM: Dulbecco’s modified Eagle’s medium
DTT: dithiothreitol
D/V: dorsal/ventral
FBS: fetal bovine serum
FP: fluorescence polarization
GO: gene ontology
HEPES: 4-(2-hydroxyethyl)-1-piperazineethanesulfonic acid
HTS: High-throughput screening
IPTG: isopropyl-D-thiogalactoside
LB: Luria-Bertani
LC: liquid chromatography
L-DOPA: L-3,4-dihydroxyphenylalanine
MOI: multiplicity of infection
MS: mass spectrometry
NTA: nickel-nitrilotriacetic acid complex
pAG-MNase: protein A-G micrococcal nuclease
PBS: phosphate-buffered saline
PDB: Protein Data Bank
pfu: plaque-forming unit
PMSF: phenylmethylsulfonyl fluoride
r.m.s.: root-mean-square
SE: standard error
SEM: standard error of the mean
TAMRA,TMR: tetramethylrhodamine
TCEP: tris(2-carboxyethyl)phosphine
TEV: tobacco etch virus
Tris: tris(hydroxymethyl)aminomethane

## ACKNOWLEDGEMENTS

GR gratefully acknowledges support from NIDA (R01DA040621; R01DA040621-03S1; R01DA040621-07S1) and the Sealy Center for Structural Biology and Molecular Biophysics (SCSB) at the University of Texas Medical Branch (UTMB) for providing research resources. JZ is partly supported by the John D. Stobo, M.D. Distinguished Chair Endowment Fund, and Edith & Robert Zinn Chair in Drug Discovery Endowment Fund. SM is funded by a Jeane B. Kempner Award (UTMB). Dr. Dan Fass, Dr. Hubert Lee, Dr. Anthony J Pastore, Galina Aglyamova, and Gregory Tan are thanked for their support with preliminary pilot studies, experimental assistance, and useful discussions. The expert assistance of Nghi D. Nguyen and useful discussions with Dr. Clifford Stephan regarding compound testing are gratefully acknowledged. Access to high-throughput instrumentation and expertise was provided by The Combinatorial Drug Discovery Program (CPRIT RP150578) at the Institute of Bioscience and Technology, Texas A&M via Dr. Clifford Stephan. We also acknowledge support from NS085171 (JC) and F30 AG085919 (CS).

## AUTHOR CONTRIBUTIONS

GR, JZ, EJN and JC conceived the experimental designs. SM, AK, and SF performed protein over-expression and protein purification, mutagenesis, all biophysical studies, and data analysis. SM, AK, SF and NT performed compound testing and analyses using biochemical and biophysical assays. YYY and MR carried out all reporter cell-based assays, compound toxicity assays, and analyses. BWH, ME, EPC, and EJN carried out the CUT&RUN and data analysis. Anil K, YL, QB, HC, and JZ carried out all chemical syntheses and quality control. SH and AJR provided critical insight and discussion of the experimental design. CS and JC carried out *in vivo* compound administration and immunohistochemical analyses. GR drafted the initial version of the manuscript, MM assisted in making figures, and all authors revised and approved it, leading to a final version.

## METHODS

### Constructs

#### Recombinant proteins overexpressed in *Sf9* insect cells

(a) full-length mouse ΔFOSB (a FOSB splice variant; UniProt ID: P13346), residues F^2^–E^237^) and (b) full-length mouse JUND (UniProt ID: J04509), residues E^2^–Y^341^. All constructs were cloned into the pFastBac1 vector and contained an N-terminal (His)_6_-tag (sequence MGHHHHHH).

#### Recombinant proteins overexpressed in E. coli

(a) mouse ΔFOSB bZIP domain (residues E^153^-K^219^); and (b) mouse JUND bZIP domain (residues Q^260^–V^326^ from UniProt J04509). Note, the amino acid sequences for these ΔFOSB bZIP and JUND bZIP domain constructs are identical between mouse and human. Both constructs were cloned into the pET21a-NESG vector and contained an N-terminal (His)₆-tag with a TEV protease cleavage site (sequence MGHHHHHHENLYFQS). Note, the JunD residue numbering is offset by 6 in the human JUND sequence (Q^266^–V^332^) compared to mouse; human numbering is used throughout this study. All plasmids were sequence-verified prior to use.

### Reagents and biological resources

#### Cell lines

AP1 luciferase reporter HEK293 cells (AP1-luc HEK293): BPS Bioscience, San Diego, CA; Cat. #60405; Neuro2A cell line: ATCC; Manassas, Virginia; catalog# CCL-131.

#### Oligonucleotides

The 19-mer *cdk5* duplex oligonucleotide (‘cdk5 oligo’), derived from the AP1 site of the cyclin-dependent kinase 5 promoter (5′-CGTCGG<u>TGACTCA</u>AAACAC-3′; AP1 site underlined), was used in various assays. The version used in fluorescence experiments (‘TMR-cdk5’) was prepared by annealing equimolar amounts of complementary strands labeled at their 5′ ends with TAMRA (Sigma-Aldrich) by heating to 95°C for 2.5 minutes followed by gradual cooling with a rate of approximately 1°C/min to room temperature. Annealed duplexes were stored at –20°C in annealing buffer (10 mM Tris, pH 8.0; 50 mM NaCl) at a concentration of 50 μM

### Chemical Synthesis

The detailed synthetic methods and experimental procedures to generate new analogs are provided as Supporting Information in the **Supplemental Material**. The chemical structures and purity of all synthesized new compounds have been characterized using ^1^H NMR, ^13^C NMR, high-resolution mass spectrometry and HPLC analyses for structural validation and quality control.

### Purification of wild-type ΔFOSB/JUND heterodimers and ΔFOSB homomers

Mouse ΔFOSB/JUND heterodimers and ΔFOSB homomers were produced in Sf9 insect cells using the Bac-to-Bac expression system (Invitrogen), following protocols previously reported (Jorissen et al, 2007; Wang et al, 2012; Kumar et al, 2022; McNeme et al, 2025). Briefly, high-titer baculoviruses (∼1 × 108 pfu/mL) were used to infect Sf9 cells at ∼1.5 × 106 cells/mL. Heterodimers were obtained by co-infection with N(His)6-ΔFOSB and N(His)6-JUND viruses at MOIs of 1.0–1.5 and 1.0–3.0, respectively. For ΔFOSB homomers, the same cell density and an MOI of 1.0–1.5 were used. Infections were carried out in 6 L SF900 cultures for 72–84 h at 28°C, shaking at 145 rpm. Cells were harvested (15 min, 3000 rpm, 4°C), resuspended in PBS, flash-frozen in liquid nitrogen, and stored at −80°C.

For protein purification, cells from 6 L cultures were resuspended in 300 mL lysis buffer (25 mM Tris pH 8.0, 0.2% [v/v] Triton X-100, 1 mM TCEP) supplemented with protease inhibitors (0.5 mM PMSF, 1 mg/mL pepstatin, 10 μg/mL leupeptin, and two tablets of cOmplete™ EDTA-free Protease Inhibitor Cocktail, Roche). After incubation on ice for 30 min, cells were lysed by sonication. The lysate was treated with 300 mM NaCl, 5 mM MgCl₂, and 50 mg/mL DNase, incubated for 1 h on ice, then supplemented with 0.5 M NaBr and 10 mM imidazole. Insoluble material was removed by centrifugation at 18,000 rpm for 30 min at 4 °C. For subsequent Ni-affinity purification, Ni-NTA resin (10 mL of a 50% slurry; Thermo Scientific), pre-equilibrated in buffer A (25 mM Tris pH 8.0, 1.0 M NaCl), was added to the clarified lysate and incubated for 3 h at 4 °C. The resin was transferred to an empty column, and bound proteins were eluted using a gradient of buffer B (25 mM Tris pH 8.0, 1.0 M NaCl, 0.5 M imidazole). Fractions containing protein were combined, diluted with 25 mM Tris pH 9.0, 1 M NaCl to a final protein concentration of 0.075 mg/mL, then dialyzed overnight against 25 mM Tris pH 9.0, 300 mM NaCl, 1 mM DTT, 0.5 mM PMSF, followed by a 3 h dialysis against 25 mM Tris pH 9.0, 75 mM NaCl, 1 mM DTT, 0.5 mM PMSF prior to anion exchange chromatography. The sample was loaded onto a Mono Q 5/50 GL column (Cytiva) equilibrated with buffer A (25 mM Tris pH 9.0, 75 mM NaCl, 1 mM DTT) and eluted with a linear 0–100% gradient of buffer B (25 mM Tris pH 9.0, 1 M NaCl, 1 mM DTT). Peak fractions were pooled, concentrated, and further purified by size-exclusion chromatography using a HiLoad 16/600 Superdex 200 16/600 size exclusion column (GE Healthcare) equilibrated in 20 mM Tris pH 8.0, 1 M NaCl. For ΔFOSB/JUND, SDS-PAGE was used to confirm co-purification of both components in a 1:1 complex. Final protein purity was assessed by SDS-PAGE. Purified ΔFOSB and ΔFOSB/JUND were flash-frozen in aliquots (2–5 mg/mL) in 20 mM Tris pH 8.0, 1 M NaCl.

### Preparation of ΔFOSB/JUND bZIP domains

ΔFOSB and JunD bZIP domains were expressed in E. coli as previously described (Yin et al, 2017, 2020; Kumar et al, 2022; McNeme et al, 2025). Briefly, 6 L cultures of E. coli Rosetta 2 (DE3) cells (Invitrogen) harboring the respective expression plasmids were grown in LB medium at 37 °C to an OD₆₀₀ of ∼0.5, cooled to 16 °C, and induced with 0.5 mM IPTG overnight. Cells were harvested by centrifugation at 4 °C for 30 min at 4000 rpm, resuspended in 10 mL PBS per 1 L of culture, flash-frozen in liquid nitrogen, and stored at −80 °C until purification.

For protein purification, cell pellets were thawed on ice, resuspended in lysis buffer (20 mM Tris pH 8.0, 250 mM NaCl, 1 mM TCEP, 0.5 mM PMSF) and treated with 1.1 mg/mL lysozyme. After a 30 min incubation on ice, cells were lysed by sonication. The lysate was supplemented with 5 mM MgCl₂, 1 M NaCl, and 30 μg/mL DNase I, and incubated for an additional 30 min on ice. Prior to Ni-affinity purification, 0.5 M NaBr and 20 mM imidazole were added, and the mixture clarified by centrifugation at 18,000 rpm for 30 min at 4 °C. Ni-NTA resin (10 mL of a 50% slurry; Thermo Scientific), pre-equilibrated in buffer A (20 mM Tris pH 8.0, 1 M NaCl, 500 mM NaBr), was added to the supernatant and incubated for 3 h at 4 °C. The resin was transferred to a column, and bound protein eluted using a gradient of buffer B (20 mM Tris pH 8.0, 1 M NaCl, 500 mM NaBr, 500 mM imidazole). To remove the (His)₆-tag, TEV protease was added to the protein-containing, combined fractions at 40 μg protease per mg target protein in digestion buffer (50 mM Tris-HCl pH 8.0, 0.5 M NaCl, 1 mM DTT, 1% (v/v) glycerol), followed by a overnight incubation at 4°C. To remove TEV protease, 1 M NaCl and 0.5 mL of Ni-NTA resin (50% slurry), equilibrated in digestion buffer, were added and the sample incubated for 1 h at room temperature. The resin was pelleted by centrifugation at 900 rpm for 10 min, the cleaved protein recovered from the supernatant, passed through a Bio-Spin® column (Bio-Rad, #7326008), and the protein concentration determined using the Bio-Rad Protein Assay (Kit I, #5000002). For heterodimer formation, ΔFOSB and JUND bZIP domains were mixed at a 1:1 molar ratio. For both ΔFOSB/JUND bZIP heterodimers and ΔFOSB bZIP homomer, buffer was then exchanged by dialysis at room temperature against 20 mM HEPES pH 7.0, 500 mM NaCl. Final preparations were concentrated to ≤5 mg/mL and purified by size exclusion chromatography at 4 °C using a Superdex HiLoad 16/600 75 pg column (Cytiva) equilibrated in GF buffer (20 mM HEPES pH 7.0, 500 mM NaCl). Purity was confirmed by SDS-PAGE. Proteins were concentrated to ∼8-10 mg/mL, flash-frozen in liquid nitrogen, and stored at −80 °C.

### Fluorescence polarization dose-response curve (FP-DRC) testing the impact of compounds on the binding of DNA to ΔFOSB/JUND and ΔFOSB

The inhibitory effect of compounds on ΔFOSB/JUND or ΔFOSB binding to DNA was evaluated in 10-point fluorescence polarization dose response curve (FP-DRC) assays following established protocols (Wang *et al*, 2012; Kumar *et al*, 2022; McNeme *et al*, 2025). Serial compound dilutions (0–200 μM) were prepared in 384-well plates (Corning #3676) using an Echo 550 Acoustic Liquid Handler (Beckman) or manually. ΔFOSB/JUND (280 nM monomer) or ΔFOSB (300–320 nM monomer) was added to wells along with the TMR-labeled ‘cdk5 oligo’ (TMR-cdk5) at a final concentration of 25 nM. Wells containing 25 nM TMR-cdk5 alone representing 100% inhibition (i.e., no protein-DNA-binding) were used as the ‘positive control’, while wells containing 25 nM TMR-cdk5 plus protein (either 280 nM ΔFOSB/JUND or 320 nM ΔFOSB) without any compound representing 0% inhibition (i.e., full protein:DNA-binding) were used as the ‘negative control’. Buffer only was used as the ‘no-protein’ control. All wells, including controls, received DMSO at a consistent final concentration of 0.5% (v/v). FP-DRC assays with the compound concentration series were run in quadruplicate for each protein complex (ΔFOSB/JUND heterodimers in 20 mM HEPES pH 7.5, 150 mM NaCl, and ΔFOSB homomers in 20 mM HEPES pH 7.5, 50 mM NaCl) and in duplicate for TMR-cdk5 alone without protein. The wells containing the compound concentration series and TMR-cdk5 (but no protein) were used to detect any possible direct interference of the compound on the oligonucleotide. Each plate included additionally 16 positive control wells (TMR-cdk5 only, 100% inhibition) and 16 negative control wells (protein + TMR-cdk5, 0% inhibition). After 15 minutes of incubation at room temperature, the FP signal was measured using either a Synergy Neo2 (BioTek) or PHERAstar (BMG LabTech) plate reader (excitation 530–540 nm, emission 590 nm). Data were fitted with GraphPad Prism 6 using a 3-parameter logistic curve (‘log(inhibition) vs. response-variable slope fixing the bottom of the plot to the averaged value of the 100%-inhibition control wells on each plate) yielding IC_50_ values, SEM values and 95% confidence intervals (CI).

### Cell-Based Reporter and Toxicity Assays in AP1-luc HEK293 cells

An AP1 reporter assay was used to assess how compounds modulate AP1-driven transcriptional activity of a luciferase reporter gene. The experimental protocol followed established methods (Kumar *et al*, 2022; McNeme *et al*, 2025). Briefly, an AP1-luciferase reporter HEK293 cell line (BPS Bioscience, USA) was cultured at 37 °C and 5% CO₂ in growth medium 1B (BPS Bioscience) supplemented with 10% fetal bovine serum (FBS; Invitrogen) and 1% penicillin/streptomycin (Hyclone). Cells were plated at a density of 6.0×10⁴ cells per well in quadruplicate across two 48-well plates. Once the cell cultures reached ∼70% confluence, the medium was switched to assay medium 1B containing 0.5% FBS to induce a 24-hour serum starvation. Compounds were added in a concentration series (0.003 −100 μM) to cells or as a vehicle control 0.5% (v/v) DMSO to cells in parallel followed by a 2-hour incubation. Subsequently, cells were exposed for 24 hours to medium containing 20% FBS (“serum-stimulated”) to induce ΔFOSB protein accumulation. After treatment, cells were lysed and the luciferase activity was quantified using the Promega luciferase assay system on a BioTek Cytation 3 plate reader. Data from two independent experiments (typically n=4) were each normalized against blank wells (i.e., vehicle control treatment), combined together and then analyzed via nonlinear regression in GraphPad Prism v10 to determine IC_50_ values with 95% confidence intervals (three parameter logistic model). To test for off-target effects, cells were incubated with compounds or with 0.5% (v/v) DMSO (vehicle control), and then exposed for 24 hours to either 1) stimulation with 20% FBS (“serum-stimulated”) conditions under which the ΔFOSB protein accumulates or 2) maintenance in serum-free medium (“non-serum-stimulated”) conditions under which negligible ΔFOSB protein accumulates (Kumar *et al*, 2022). In parallel, cell viability assays were performed under identical conditions to the AP1 reporter assay to evaluate compound-induced toxicity. After a two-hour incubation with compound or vehicle (0.5% (v/v) DMSO), the medium was exchange for that containing 20% FBS (“serum-stimulated”) followed by 24 hours incubation. Cell viability was determined using the CellTiter-Glo luminescent assay (Promega) according to the manufacturer’s protocol, and luminescence was measured using a BioTek Cytation 3 reader. Statistical analyses were performed in GraphPad Prism v.10.0 (GraphPad Software), and data are presented as means ± standard error of the mean (SEM).

### Mouse liver microsomal stability assays

Compound stability was assessed in mouse liver microsomes (238.5 ml; 0.63 mg/mL) incubated with 1.5 ml compound (0.2 mM stock in 20% DMSO) at 37 °C in buffer containing 50 mM potassium phosphate (pH 7.4) in the presence or absence of 1 mM NADPH by Biodura-Sundia. The final volume per sample was 300 ml containing end concentration of liver microsomes (0.5 mg/mL), 1 μM compound, and final DMSO concentration 0.125% (v/v). Samples were collected at 0, 5, 15, 30, and 60 minutes (with NADPH) and at 0 and 60 minutes (without NADPH), and the reaction was quenched by the addition of acetonitrile. After centrifugation, an internal standard was added to the supernatants, which were analyzed by mass spectrometry (LS/MS/MS). Compound disappearance over time was used to calculate half-life (t½, in minutes), *in vitro* intrinsic clearance (CL_int_, in μl/min/mg protein), and predicted hepatic clearance (CL_hep_ in μl/min/kg).

### Administration of YL0441 in vivo

For in vivo studies, 2–3-month-old heterozygous transgenic male and female mice expressing human amyloid precursor protein (APP) carrying Swedish (K670N, M671L) and Indiana (V717F) mutations linked to familial Alzheimer’s disease (Line J20; MMRRC_034836-JAX; hAPP770 transgene) were used (Mucke et al, 2000). Using a cannula connected to an Alzet micro-osmotic pump (model 1003D), YL0441 was delivered unilaterally into the right dorsal hippocampus. Micro-osmotic pumps were each filled with vehicle (0.02% DMSO/0.9% saline) or YL0441 (50 μM compound in 0.2% DMSO/0.9% saline) per manufacturer’s instructions. The brain infusion cannula was connected to the pump using medical-grade polyvinyl chloride catheter tubing and primed overnight in 0.9% sterile saline before implantation. To permit the unilateral infusion of compound or vehicle, one filled micro-osmotic pump per mouse was implanted subcutaneously in the intrascapular region, while the tip of the cannula targeted stereotactic coordinates: dorsal-ventral (D/V), –2.0 mm; anterior-posterior (A/P), –2.1 mm; and medial-lateral (M/L), –1.2 mm, and the cannula base was affixed to the skull using Superglue. The lot of Alzet 1003D micro-osmotic pumps used delivered fluid at a rate of 0.96 μL/h yielding a final drug delivery of 18.5 ng YL0441/h. YL0441 or vehicle was infused for three days into the hippocampus. Mice were then anesthetized with commercial euthanasia solution, trans-cardially perfused with ice-cold 0.9% saline solution, and the brains collected and hemisected. The left hemibrain was fixed in 4% paraformaldehyde and used for immunohistochemistry. The right hemibrain was flash-frozen on dry ice, and then later the hippocampus was isolated, and the dorsal two-thirds sub-dissected in order to use for CUT&RUN analysis. All procedures were approved by the Institutional Animal Care and Use Committees of Baylor College of Medicine. Before sectioning on a freezing sliding microtome, the fixed hemibrains were cryoprotected in 30% sucrose in phosphate-buffered saline. Subsequently, serial coronal sections (30 μm thickness) were divided into 10 subseries, each containing every tenth section throughout the rostral-caudal extent of the brain. One subseries was used to stain for ΔFOSB applying the avidin-biotin diaminobenzidine (DAB) method, with as primary antibody, the rabbit anti-ΔFOSB antibody (1:5000; Cell Signaling, D3S8R), and as secondary antibody, the biotinylated goat anti-rabbit (1:200; Vector, BA-1000), with DAB as the chromogen. All samples were processed, stained, and imaged at the same time, using identical parameters and imaging settings. To quantify ΔFOSB protein levels in the tissue, an experimenter blinded to treatment evaluated and confirmed ΔFOSB induction in the transgenic mice. The ImageJ software was used to measure the mean gray value of the dentate gyrus granule cell layer, which was then normalized to the mean gray value of the stratum radiatum and averaged over two consecutive sections for each sample. For these studies, 11 APP mice (6 male and 5 female) were treated with YL0441, and 10 mice (6 male and 4 female) were treated with vehicle.

### CUT&RUN-sequencing

#### Nuclei Isolation

Nuclei were isolated from dorsal hippocampal tissue from APP mice which had been infused with YL0441 or vehicle. Tissue was homogenized in lysis buffer (320 mM sucrose, 5 mM CaCl_2_, 0.1 mM EDTA, 10 mM Tris-HCl, pH 8.0, 1 mM DTT, 0.1% Triton X-100, 1.5 mM spermidine, and protease inhibitors) using a Dounce homogenizer (30-40 strokes per pestle). Homogenates were then passed through a 40 μm strainer, transferred to ultracentrifuge tubes, and a sucrose cushion layered underneath (1.8 M sucrose, 10 mM Tris-HCl, pH 8.0, 1 mM DTT, 1.5 mM spermidine, and protease inhibitors), upon which the nuclei were then pelleted by ultracentrifugation at 24,000 rpm for 1.25 h at 4°C.

#### CUT&RUN Assay

CUT&RUN was performed as described previously (Yeh *et al*, 2023; McNeme *et al*, 2025; Lynch *et al*, 2025). Briefly nuclei were resuspended in wash buffer (20 mM HEPES-NaOH, pH 7.5, 150 mM NaCl, 0.5 mM spermidine, 0.1% Triton X-100, 0.1% Tween-20, 0.1% BSA, and protease inhibitors) and then incubated on BioMag®Plus Concanavalin A beads (BP531, Bang Laboratories) for 10 min at room temperature in binding buffer (20 mM HEPES-KOH, pH 7.9, 10 mM KCl, 1 mM CaCl_2_, and 1 mM MnCl_2_). Beads were magnetically retrieved and then incubated with primary anti-ΔFOSB antibody (Cell Signaling, #D3S8R) (1:50) or IgG control (1:50) overnight at 4°C in wash buffer. The beads were washed in buffer, and then incubated with pAG-MNase (EpiCypher) for 1 h at 4°C. The beads were then washed again in low-salt buffer, incubated in digestion buffer (3.5 mM HEPES-NaOH, pH 7.5, 10 mM CaCl_2_, 0.1% Triton X-100, and 0.1% Tween-20) for 5 min on a pre-chilled 4 °C block, and then the pAG-MNase digestion was stopped with ED:ET STOP buffer (170 mM NaCl, 20 mM EDTA, 4 mM EGTA, 0.1% Triton X-100, 0.1% Tween-20, 25 μg/mL RNase, and 20 μg/mL glycogen). The DNA fragments were eluted by shaking at 37°C for 30 min, centrifuged at 16,000 RCF for 5 min at 4°C, and then precipitated with ethanol. Three-six independent libraries were generated per condition, using the NEB Next® Ultra™ II DNA Library Prep Kit (New England BioLabs) and sequenced on an Illumina HiSeq 4000 at a minimum depth of ≥30 million paired-end reads per sample using 2 × 150 bp paired-end configuration and analyzed with SEACR (relaxed threshold).

#### Data Analysis

FASTQ files were processed with the NGS-Data-Charmer pipeline (https://github.com/shenlab-sinai/NGS-Data-Charmer). Trim-Galore (v0.6.5) was used for adapter trimming, followed by secondary trimming with Cutadapt (v2.10). Reads were aligned to the mm10 genome using HISAT2 (v2.2.0), and duplicate reads were removed using the Picard (v3.0) ‘MarkDuplicates’ module. For visualization purposes, de-duplicated reads were converted to bigwig files with the Deeptools (v3.5.0) ‘bamCoverage’ module (--binSize 10 --normalizeUsing RPKM), then visualized as tracks using IGVtools (v2.5.3). MACS2 (v2.2.6) (-f BAMPE -q 0.01 --keep-dup) was used for peak calling after subtracting the IgG samples pooled from each APP mouse as background. Peak files were annotated using ChIPseeker (v1.22.1), and heatmaps were created for each group. Differential peak correlations were visualized using DiffBind (v3.9) with p < 0.0001, merged, and filtered at an FDR of 10%. Bigwig files were visualized using IGV (v2.12.2). No differences were seen between male and female mice, hence, all analyses combined the two sexes. The Homer software was used to perform Known Motif Enrichment analyses and motif discovery (v4.9) (Heinz *et al*, 2010). The nuclei from mice from Cohort 1 and Cohort 2 were processed, and sequenced separated, and the data combined for final bioinformatic analyses.

### Data and software availability

The CUT&RUN data sets are under submission to NIH GEO.

