## Supplemental Material for "Efficient *in vivo* pharmacological inhibition of ΔFOSB, an AP1 transcription factor, in brain"

**Supporting Information:**

**Synthetic routes and experimental procedures to generate new analogs for ΔFOSB inhibitors**

**Experimental Section**

**General.** All chemicals and solvents were obtained from commercial suppliers and used without further purification unless specified. Analytical TLC was performed on silica gel 60F 254 plates (Merck) with UV detection at 254 nm. Preparative column chromatography was carried out on silica gel 60 (70−230 mesh, flash). NMR spectra were recorded on a Bruker spectrometer (300 MHz for ^1^H and 75 MHz for ^13^C ) in *CDCl_3_* or *methanol-d_4_* or *DMSO-d_6_*. Chemical shifts (*δ*) are reported in ppm, with TMS (δ = 0 ppm) or residual solvent signals (CDCl_3_: δ = 7.265 ppm, DMSO: δ = 2.5 ppm) as internal references for ^1^H NMR, and CDCl_3_ (δ = 77.16 ppm) or DMSO (δ = 39.51 ppm) for ^13^C NMR. Coupling constants are reported in Hz. High-resolution mass spectra (HRMS) were obtained on a Thermo Fisher LTQ Orbitrap Elite mass spectrometer. Parameters include the following: the nano ESI spray voltage was 1.8 kV, capillary temperature was 275 °C, and the resolution was 60,000; ionization was achieved by positive mode. The purity of final compounds was determined by HPLC on a Shimadzu system (model: CBM-20ALC-20ADSPD-20AUV/vis) using a Waters μBondapak C18 column (300 × 3.9 mm), with a flow rate of 0.5 mL/min, UV detection at 270 and 254 nm. A linear gradient from 10% acetonitrile in water (0.1% TFA) to 100% acetonitrile (0.1% TFA) over 20 min, followed by 30 min with the final solvent, was used.

**Synthesis of acrylamide-derived aryl sulfonic acids:**

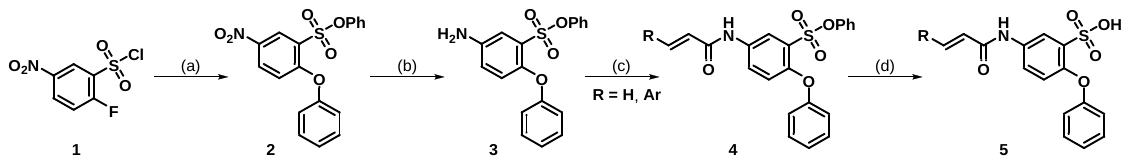

(a) phenol (3.0 equiv.), NaH (3.0 equiv.), DCM (0.25 M), 60 ^o^C, 2 h, 89% yield; (b) Pd/C (10 mol%), H_2_, DCM (0.30 M), 25 ^o^C, 24 h, 91% yield; (c) Acrylic acid derivatives (2.0 equiv.), PPAA (2.0 equiv.), TEA (2.0 equiv.), DCM (0.2 M), rt, 6 h, quantitative yield; (d) (i) aq. NaOH (2.0 M, 5.0 equiv.), dioxane:water (v/v 3:1, 0.1 M), 100 ^o^C, 6 h. (ii) aq. HCl (2.0 N, pH adjusted to approximately 3~4), 22-77% yield.

***Phenyl 5-nitro-2-phenoxybenzenesulfonate (2, AK0203)***

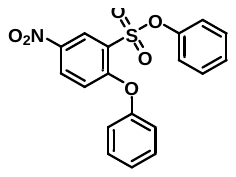

An oven-dried 200 mL single-neck round-bottom flask equipped with a magnetic stir bar was charged with 2-fluoro-5-nitrobenzenesulfonyl chloride (**1**) (4.8 g, 20.0 mmol, 1.0 equiv.) and phenol (2.8 g, 30.0 mmol, 3.0 equiv.) in dichloromethane (DCM, 80 mL, 0.25 M). Sodium hydride (NaH, 60% dispersion in mineral oil) (1.2 g, 30.0 mmol, 3.0 equiv.) was then slowly added to the reaction mixture. The reaction mixture was refluxed at 60 °C for 2 hours. After completion, the mixture was cooled to room temperature and quenched with 50 mL of ice water. The aqueous layer was extracted with DCM (3 × 75 mL), and the combined organic extracts were dried over anhydrous sodium sulfate. The solvent was removed under reduced pressure, and the crude product was purified by column chromatography on silica gel (70–230 mesh) using a gradient of ethyl acetate/hexane (5:95 to 10:90 v/v) as eluent to afford the desired product, phenyl 5-nitro-2-phenoxybentofzenesulfonate (**2**, **AK0203**), as a yellow solid (6.6 g, 89% yield). ^1^H NMR (300 MHz, CDCl_3_) δ 8.80 (t, *J* = 2.9 Hz, 1H), 8.35 (dd, *J* = 9.2, 2.9 Hz, 1H), 7.56 – 7.41 (m, 2H), 7.39 – 7.25 (m, 4H), 7.24 – 7.11 (m, 4H), 7.03 – 6.90 (m, 1H). ^13^C NMR (75 MHz, CDCl_3_) δ 161.2, 153.6, 149.5, 141.6, 131.0, 130.8, 130.1, 128.1, 127.7, 126.8, 125.8, 122.0, 120.9, 117.3.

***Phenyl 5-amino-2-phenoxybenzenesulfonate (3, AK0205)***

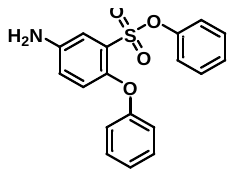

An oven-dried 100 mL single-neck round-bottom flask equipped with a magnetic stir bar was charged with phenyl 5-nitro-2-phenoxybenzenesulfonate (**2**, **AK0203**) (5.6 g, 15.0 mmol, 1.0 equiv.) in dichloromethane (DCM, 50 mL, 0.3 M). Palladium on carbon (Pd/C, 10 wt.% on activated carbon) (160 mg, 10 mol%) was added at room temperature. The flask was evacuated and refilled with a hydrogen balloon. The reaction mixture was stirred at room temperature for 24 hours. After completion of the reaction (monitored by TLC), the mixture was filtered through a bed of Celite. The filtrate was concentrated under reduced pressure, and the crude product was purified by column chromatography on silica gel (70–230 mesh) using a gradient of methanol/dichloromethane (1:99 to 3:97 v/v) as eluent to afford the desired product, phenyl 5-amino-2-phenoxybenzenesulfonate (**3**, **AK0205**), as a light-yellow solid (4.6 g, 91% yield). ^1^H NMR (300 MHz, CDCl_3_) δ 7.40 – 6.94 (m, 11H), 6.83 (d, *J* = 10.4 Hz, 2H), 3.76 (s, 2H). ^13^C NMR (75 MHz, CDCl_3_) δ 157.3, 149.7, 146.8, 142.6, 129.8, 129.7, 127.2, 127.0, 123.6, 122.2, 122.0, 121.9, 118.5, 116.4.

***A. The general procedure for the amide coupling reaction***

An oven-dried 50 mL screw-cap reaction vial equipped with a magnetic stir bar was charged with the acrylic acid derivatives (2.0 equiv.) and polyphosphoric acid anhydride (PPAA, 50% in ethyl acetate) (2.0 equiv.) in dichloromethane (DCM, 0.2 M). The reaction mixture was stirred at room temperature for 30 minutes. Subsequently, phenyl 5-amino-2-phenoxybenzenesulfonate (**3**, **AK0205**) (1.0 equiv.) and triethylamine (2.0 equiv.) were added slowly to the reaction mixture. Stirring was continued at room temperature for 12 hours. Upon completion, the reaction was quenched with 10 mL of ice water and extracted with DCM (3 × 25 mL). The combined organic layers were washed with aqueous NaOH (1.0 M, 20 mL), dried over anhydrous sodium sulfate, and concentrated under reduced pressure to afford the desired amide coupling product (**4**), which was used directly in the subsequent step without further purification.

***B. The general procedure for the cleavage of phenyl group from sulfonate esters***

An oven-dried 25 mL screw-cap reaction vial equipped with a magnetic stir bar was charged with the amide coupling product (**4**) (1.0 equiv.) and aq. NaOH (2.0 M, 5.0 equiv.) in a dioxane:water (v/v 3:1, 0.1 M) mixture. The vial was sealed with a screw cap and placed in a pre-heated metal block at 100 °C for 6 hours. Upon completion of the reaction, the mixture was cooled to room temperature and acidified with 2 N aqueous HCl to adjust the pH to approximately 3~4. The resulting mixture was extracted with ethyl acetate (3 × 25 mL), and the combined organic layers were dried over anhydrous sodium sulfate. The solvent was removed under reduced pressure, and the crude product was purified by column chromatography on silica gel (70–230 mesh), using a gradient of methanol/dichloromethane (5:95 to 10:90 v/v) as the eluent, to afford the desired sulfonic acid products (**5**).

***5-(4-Bromobenzamido)-2-phenoxybenzenesulfonic acid (YL0440)***

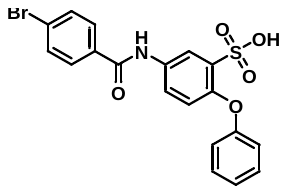

Prepared as shown in *the general experimental procedures B* on a 0.20 mmol scale. Yield = 62% (55 mg); Appearance – Yellow Solid; ^1^H NMR (300 MHz, methanol-*d*) δ 8.16 (d, *J* = 2.7 Hz, 1H), 7.95 – 7.80 (m, 3H), 7.75 – 7.66 (m, 2H), 7.42 – 7.31 (m, 2H), 7.21 – 7.07 (m, 3H), 6.87 (d, *J* = 8.9 Hz, 1H). ^13^C NMR (75 MHz, methanol-*d*) δ 166.4, 157.3, 151.5, 135.4, 133.7, 132.8, 131.5, 129.3, 129.1, 126.0, 124.6, 123.3, 121.7, 119.4, 119.2. MS (ESI) m/z [M + H]^+^ Calculated for C_19_H_15_BrNO_5_S 447.9854; found 447.9858.

***(E)-5-(3-(furan-2-yl)acrylamido)-2-phenoxybenzenesulfonic acid (YL0441)***

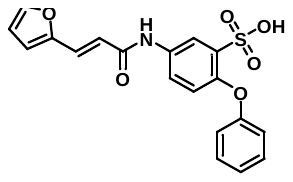

Prepared as shown in *the general experimental procedures B* on a 0.20 mmol scale. Yield = 55% (45 mg); Appearance – Yellow-brown Solid; ^1^H NMR (300 MHz, methanol-*d_4_*) δ 8.15 (d, *J* = 2.7 Hz, 1H), 7.83 (dd, *J* = 8.9, 2.7 Hz, 1H), 7.63 (d, *J* = 1.8 Hz, 1H), 7.45 (d, *J* = 15.4 Hz, 1H), 7.36 (t, *J* = 7.9 Hz, 2H), 7.12 (tt, *J* = 8.5, 1.1 Hz, 3H), 6.83 (d, *J* = 8.9 Hz, 1H), 6.74 (d, *J* = 3.4 Hz, 1H), 6.68 – 6.50 (m, 2H). ^13^C NMR (75 MHz, methanol-*d_4_*) δ 165.3, 157.3, 151.3, 151.0, 144.6, 135.4, 133.3, 129.3, 128.2, 123.4, 123.2, 120.5, 119.4, 119.3, 118.1, 114.2, 112.1. MS (ESI) m/z [M + H]^+^ Calculated for C_19_H_15_NO_6_S 386.0698; found 386.0686.

***5-Cinnamamido-2-phenoxybenzenesulfonic acid (QB0280)***

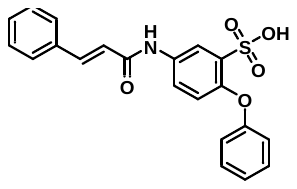

Prepared as shown in *the general experimental procedures B* on a 0.15 mmol scale. Yield = 69% (39 mg); Appearance – White Solid; ^1^H NMR (300 MHz, DMSO-*d*_6_) δ 10.28 (s, 1H), 7.99 (d, *J* = 2.7 Hz, 1H), 7.85 (dd, *J* = 8.7, 2.7 Hz, 1H), 7.69 – 7.50 (m, 3H), 7.49 – 7.20 (m, 5H), 7.09 – 6.72 (m, 5H). ^13^C NMR (75 MHz, DMSO-*d*_6_) δ 165.6, 157.6, 151.8, 139.3, 136.5, 136.4, 130.2, 129.9, 129.8, 129.0, 128.4, 124.4, 122.1, 120.6, 117.8, 115.1. MS (ESI) m/z [M + H]^+^ Calculated for C_21_H_17_NO_5_S 396.0906; found 396.1012.

***(E)-5-(3-(4-Methoxyphenyl)acrylamido)-2-phenoxybenzenesulfonic acid (QB0292)***

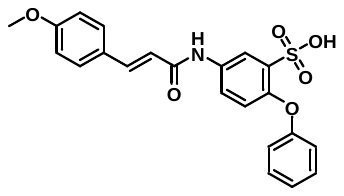

Prepared as shown in *the general experimental procedures B* on a 0.16 mmol scale. Yield = 57% (39 mg); Appearance – White Solid; ^1^H NMR (300 MHz, DMSO-*d*_6_) δ 10.19 (s, 1H), 7.97 (d, *J* = 2.7 Hz, 1H), 7.84 (dd, *J* = 8.7, 2.7 Hz, 1H), 7.62 – 7.47 (m, 3H), 7.30 (dd, *J* = 8.6, 7.3 Hz, 2H), 7.02 (dd, *J* = 8.1, 6.3 Hz, 3H), 6.89 (dt, *J* = 7.9, 1.1 Hz, 2H), 6.78 (d, *J* = 8.8 Hz, 1H), 6.68 (d, *J* = 15.7 Hz, 1H), 3.80 (s, 3H), 3.17 (d, *J* = 4.8 Hz, 2H). ^13^C NMR (75 MHz, DMSO-*d*_6_) δ 165.6, 160.8, 157.6, 151.8, 142.0, 136.5, 130.2, 129.9, 129.6, 128.4, 124.5, 122.1, 120.6, 117.8, 115.1, 114.7, 55.3. MS (ESI) m/z [M + H]^+^ Calculated for C_22_H_19_NO_6_S 426.1011; found 426.1014.

***(E)-5-(3-(3,4-Dimethoxyphenyl)acrylamido)-2-phenoxybenzenesulfonic acid (QB0293)***

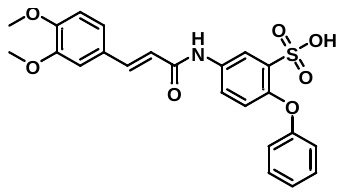

Prepared as shown in *the general experimental procedures B* on a 0.15 mmol scale. Yield = 61% (43 mg); Appearance – White Solid; ^1^H NMR (300 MHz, DMSO-*d*_6_) δ 10.19 (s, 1H), 7.98 (d, *J* = 2.7 Hz, 1H), 7.84 (dd, *J* = 8.7, 2.7 Hz, 1H), 7.51 (d, *J* = 15.6 Hz, 1H), 7.34 – 7.26 (m, 2H), 7.24 – 7.14 (m, 2H), 7.01 (dd, *J* = 7.8, 4.8 Hz, 2H), 6.93 – 6.61 (m, 5H), 3.81 (d, *J* = 6.4 Hz, 6H). ^13^C NMR (75 MHz, DMSO-*d*_6_) δ 165.6, 157.6, 151.8, 150.5, 150.0, 142.5, 136.5, 130.2, 129.9, 127.8, 124.5, 122.2, 122.1, 120.6, 118.7, 117.8, 115.1, 112.3, 110.4, 56.0, 55.9. MS (ESI) m/z [M + H]^+^ Calculated for C_23_H_21_NO_7_S 456.1117; found 456.1124.

***(E)-5-(3-(4-Chlorophenyl)acrylamido)-2-phenoxybenzenesulfonic acid (QB0294)***

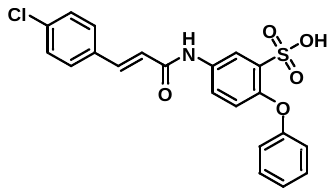

Prepared as shown in *the general experimental procedures B* on a 0.15 mmol scale. Yield = 77% (49 mg); Appearance – White Solid; ^1^H NMR (300 MHz, DMSO-*d*_6_) δ 10.30 (s, 1H), 7.98 (d, *J* = 2.7 Hz, 1H), 7.85 (dd, *J* = 8.7, 2.7 Hz, 1H), 7.70 – 7.45 (m, 5H), 7.30 (dd, *J* = 8.6, 7.3 Hz, 2H), 7.02 (t, *J* = 7.3 Hz, 1H), 6.93 – 6.71 (m, 4H). ^13^C NMR (75 MHz, DMSO-*d*_6_) δ 165.6, 157.61, 151.8, 142.0, 136.5, 134.5, 134.4, 130.2, 129.9, 129.2, 129.0, 124.5, 122.1, 120.6, 117.8, 115.1. MS (ESI) m/z [M + H]^+^ Calculated for C_21_H_16_ClNO_5_S 430.0516; found 432.0504.

***(E)-5-(3-(3,4-Dichlorophenyl)acrylamido)-2-phenoxybenzenesulfonic acid (QB0295)***

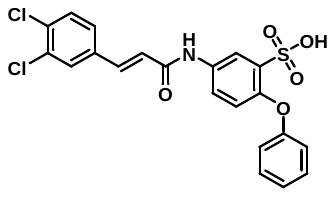

Prepared as shown in *the general experimental procedures B* on a 0.13 mmol scale. Yield = 67% (40 mg); Appearance – White Solid; ^1^H NMR (300 MHz, DMSO-*d*_6_) δ 10.31 (s, 1H), 7.98 (d, *J* = 2.7 Hz, 1H), 7.92 – 7.79 (m, 2H), 7.71 (d, *J* = 8.3 Hz, 1H), 7.63 (dd, *J* = 8.5, 2.0 Hz, 1H), 7.56 (d, *J* = 15.8 Hz, 1H), 7.30 (dd, *J* = 8.6, 7.3 Hz, 2H), 7.02 (t, *J* = 7.4 Hz, 1H), 6.93 – 6.72 (m, 4H). ^13^C NMR (75 MHz, DMSO-*d*_6_) δ 165.6, 157.6, 151.8, 140.7, 136.5, 135.2, 134.3, 133.9, 130.9, 130.2, 130.2, 129.9, 128.2, 124.5, 122.1, 120.6, 118.7, 117.8, 115.1. MS (ESI) m/z [M + H]^+^ Calculated for C_21_H_15_Cl_2_NO_5_S 464.0126; found 464.0138.

***(E)-5-(3-(4-Fluorophenyl)acrylamido)-2-phenoxybenzenesulfonic acid (QB0301)***

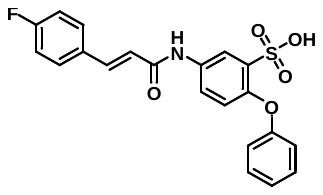

Prepared as shown in *the general experimental procedures B* on a 0.20 mmol scale. Yield = 44% (37 mg); Appearance – White Solid; ^1^H NMR (300 MHz, DMSO-*d*_6_) δ 10.27 (s, 1H), 7.98 (d, *J* = 2.7 Hz, 1H), 7.84 (dd, *J* = 8.7, 2.8 Hz, 1H), 7.69 (dd, *J* = 8.7, 5.6 Hz, 2H), 7.58 (d, *J* = 15.8 Hz, 1H), 7.38 – 7.21 (m, 4H), 7.02 (t, *J* = 7.3 Hz, 1H), 6.93 – 6.84 (m, 2H), 6.82 – 6.71 (m, 2H). ^13^C NMR (75 MHz, DMSO-*d*_6_) δ 165.6, 157.6, 151.8, 142.0, 136.5, 131.2, 130.2, 129.9, 129.1, 129.1, 124.5, 122.1, 120.6, 117.8, 115.1, 114.8, 114.6. MS (ESI) m/z [M + H]^+^ Calculated for C_21_H_16_FNO_5_S 414.0811; found 414.0816.

***(******E)-2-Phenoxy-5-(3-(4-(trifluoromethyl)phenyl)acrylamido)benzenesulfonic acid (QB0309)***

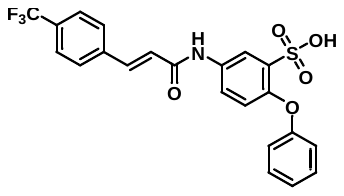

Prepared as shown in *the general experimental procedures B* on a 0.16 mmol scale. Yield = 49% (36 mg); Appearance – White Solid; ^1^H NMR (300 MHz, DMSO-*d*_6_) δ 8.76 (d, *J* = 2.2 Hz, 1H), 8.53 (dd, *J* = 4.8, 1.6 Hz, 1H), 8.06 – 7.87 (m, 2H), 7.76 (dd, *J* = 8.7, 2.7 Hz, 1H), 7.52 – 7.37 (m, 2H), 7.33 – 7.22 (m, 2H), 7.05 – 6.82 (m, 4H), 6.71 (d, *J* = 8.7 Hz, 1H). ^13^C NMR (75 MHz, DMSO-*d*_6_) δ 165.6, 157.6, 151.8, 142.0, 136.5, 133.0, 130.6, 130.4, 130.2, 129.9, 128.3, 128.3, 125.7, 125.7, 125.7, 125.6, 125.0, 124.5, 122.1, 120.6, 117.8, 115.1. MS (ESI) m/z [M + H]^+^ Calculated for C_22_H_16_F_3_NO_5_S 464.0780; found 464.0789.

***(E)-5-(3-(3,4-Dihydroxyphenyl)acrylamido)-2-phenoxybenzenesulfonic acid (QB0311)***

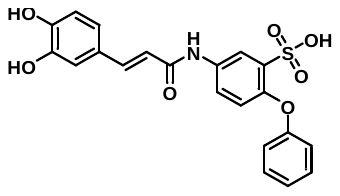

Prepared as shown in *the general experimental procedures B* on a 0.16 mmol scale. Yield = 22% (15 mg); Appearance – White Solid; ^1^H NMR (300 MHz, DMSO-*d*_6_) δ 10.14 (s, 1H), 7.95 (d, *J* = 2.6 Hz, 1H), 7.84 (dd, *J* = 8.7, 2.7 Hz, 1H), 7.38 (d, *J* = 15.7 Hz, 1H), 7.33 – 7.26 (m, 2H), 7.01 (s, 2H), 6.89 (d, *J* = 8.2 Hz, 4H), 6.77 (d, *J* = 8.7 Hz, 2H), 6.53 (d, *J* = 15.3 Hz, 1H), 1.81 (s, 1H), 1.24 (s, 1H). ^13^C NMR (75 MHz, DMSO-*d*_6_) δ 165.6, 157.6, 151.8, 148.3, 147.1, 142.5, 136.5, 130.2, 129.9, 128.0, 124.5, 122.4, 122.1, 120.6, 118.7, 117.8, 116.3, 115.1, 115.1. MS (ESI) m/z [M + H]^+^ Calculated for C_21_H_17_NO_7_S 428.0804; found 428.0810.

***(E)-5-(3-(4-Nitrophenyl)acrylamido)-2-phenoxybenzenesulfonic acid (QB0323)***

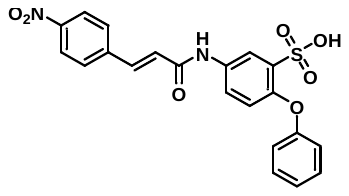

Prepared as shown in *the general experimental procedures B* on a 0.30 mmol scale. Yield = 22% (16 mg); Appearance – Dark-brown Solid; ^1^H NMR (300 MHz, DMSO-*d*_6_) δ 10.43 (s, 1H), 8.29 (d, *J* = 8.7 Hz, 2H), 8.00 (d, *J* = 2.7 Hz, 1H), 7.94 – 7.82 (m, 3H), 7.69 (d, *J* = 15.8 Hz, 1H), 7.35 – 7.27 (m, 2H), 7.07 – 6.75 (m, 6H). ^13^C NMR (75 MHz, DMSO-*d*_6_) δ 165.1, 149.6, 149.5, 149.4, 146.2, 145.9, 144.8, 144.8, 144.7, 142.7, 137.3, 124.0, 121.8, 114.4, 113.9, 113.3, 112.2, 108.8, 108.7, 108.7. MS (ESI) m/z [M + H]^+^ Calculated for C_21_H_16_N_2_O_7_S 441.0756; found 441.0759.

***(E)-5-(3-(4-(Dimethylamino)phenyl)acrylamido)-2-phenoxybenzenesulfonic acid(QB0326)***

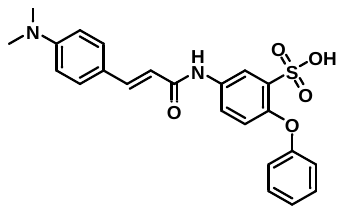

Prepared as shown in *the general experimental procedures B* on a 0.30 mmol scale. Yield = 26% (36 mg); Appearance – White Solid; ^1^H NMR (300 MHz, DMSO-*d*_6_) δ 10.08 (s, 1H), 7.96 (d, *J* = 2.7 Hz, 1H), 7.84 (dd, *J* = 8.8, 2.7 Hz, 1H), 7.55 – 7.37 (m, 3H), 7.36 – 7.23 (m, 2H), 7.10 – 6.96 (m, 1H), 6.94 – 6.71 (m, 5H), 6.57 (d, *J* = 15.7 Hz, 1H), 2.99 (s, 6H). ^13^C NMR (75 MHz, DMSO-*d*_6_) δ 164.5, 159.0, 151.3, 148.9, 140.8, 140.4, 129.7, 129.6, 122.4, 121.4, 121.2, 120.2, 118.7, 117.4, 117.4, 117.3, 113.0, 39.5, 39.3. MS (ESI) m/z [M + H]^+^ Calculated for C_23_H_22_N_2_O_5_S 439.1328; found 439.1339.

***(E)-5-(3-(4-Aminophenyl)acrylamido)-2-phenoxybenzenesulfonic acid (QB0343)***

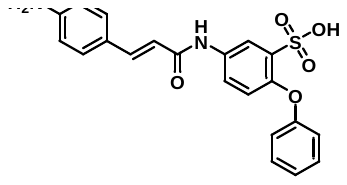

Prepared as shown in the *general experimental procedures D* on a 0.10 mmol scale. Yield = 92% (37 mg); Appearance – White Solid; ^1^H NMR (300 MHz, DMSO-*d*_6_) δ 10.43 (s, 1H), 8.29 (d, *J* = 8.7 Hz, 2H), 8.00 (d, *J* = 2.7 Hz, 1H), 7.94 – 7.82 (m, 3H), 7.69 (d, *J* = 15.8 Hz, 1H), 7.35 – 7.27 (m, 2H), 7.07 – 6.75 (m, 6H). ^13^C NMR (75 MHz, DMSO-*d*_6_) δ 166.5, 157.2, 152.6, 149.2, 147.8, 141.5, 133.5, 129.9, 128.0, 124.5, 122.4, 120.6, 117.8, 114.8, 114.1, 112.3. MS (ESI) m/z [M + H]^+^ Calculated for C_21_H_18_N_2_O_5_S 411.1015; found 411.1009.

***5-(3-([1,1'-Biphenyl]-4-yl)acrylamido)-2-phenoxybenzenesulfonic acid (QB0348)***

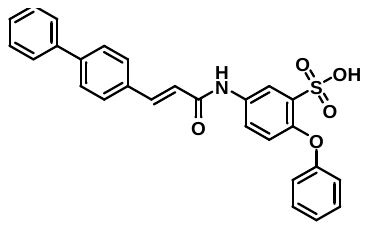

Prepared as shown in *the general experimental procedures B* on a 0.16 mmol scale. Yield = 56% (44 mg); Appearance – White Solid; ^1^H NMR (300 MHz, DMSO-*d*_6_) δ 10.22 (s, 1H), 7.93 (d, *J* = 2.8 Hz, 1H), 7.70 (dd, *J* = 8.8, 2.9 Hz, 1H), 7.66 – 7.53 (m, 6H), 7.49 (d, *J* = 15.7 Hz, 1H), 7.39 – 7.30 (m, 2H), 7.25 (dd, *J* = 8.4, 6.0 Hz, 1H), 7.18 (t, *J* = 7.9 Hz, 2H), 6.90 (t, *J* = 7.3 Hz, 1H), 6.83 – 6.71 (m, 3H), 6.67 (d, *J* = 8.8 Hz, 1H). ^13^C NMR (75 MHz, DMSO-*d*_6_) δ 163.9, 158.8, 149.4, 141.7, 140.0, 139.9, 139.7, 134.6, 134.3, 129.8, 129.5, 128.8, 128.3, 127.6, 127.1, 122.7, 121.6, 121.3, 120.4, 118.8. MS (ESI) m/z [M + H]^+^ Calculated for C_27_H_21_NO_5_S 472.1219; found 472.1224.

***(E)-2-Phenoxy-5-(3-(pyridin-4-yl)acrylamido)benzenesulfonic acid (QB0310)***

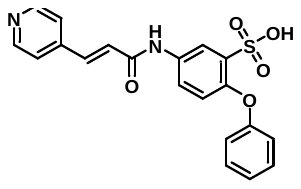

Prepared as shown in *the general experimental procedures B* on a 0.20 mmol scale. Yield = 73% (57 mg); Appearance – Light-yellow Solid; ^1^H NMR (300 MHz, DMSO-*d*_6_) δ 10.38 (s, 1H), 8.00 (d, *J* = 2.8 Hz, 1H), 7.89 – 7.76 (m, 5H), 7.65 (d, *J* = 15.7 Hz, 1H), 7.37 – 7.23 (m, 2H), 7.09 – 6.85 (m, 4H), 6.80 (d, *J* = 8.7 Hz, 1H). ^13^C NMR (75 MHz, DMSO-*d*_6_) δ 164.3, 150.1, 149.5, 148.1, 140.0, 134.6, 134.2, 131.8, 129.6, 128.4, 124.4, 122.5, 122.2, 121.5, 121.1, 118.6. MS (ESI) m/z [M + H]^+^ Calculated for C_20_H_16_NO_5_S 397.0858; found 397.1861.

***(E)-2-Phenoxy-5-(3-(pyridin-2-yl)acrylamido)benzenesulfonic acid (QB0324)***

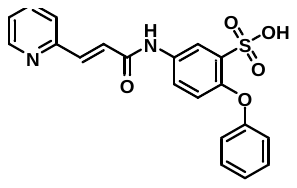

Prepared as shown in *the general experimental procedures B* on a 0.20 mmol scale. Yield = 68% (54 mg); Appearance – Light-yellow Solid; ^1^H NMR (300 MHz, DMSO-*d*_6_) δ 10.58 (s, 1H), 8.78 (dd, *J* = 5.4, 1.7 Hz, 1H), 8.21 (td, *J* = 7.8, 1.7 Hz, 1H), 8.08 (d, *J* = 2.7 Hz, 1H), 7.97 (d, *J* = 7.9 Hz, 1H), 7.83 (dd, *J* = 8.8, 2.7 Hz, 1H), 7.74 – 7.60 (m, 2H), 7.41 – 7.24 (m, 3H), 7.02 (td, *J* = 7.3, 1.2 Hz, 1H), 6.92 – 6.85 (m, 2H), 6.81 (d, *J* = 8.7 Hz, 1H). ^13^C NMR (75 MHz, DMSO-*d*_6_) δ 162.7, 158.8, 151.0, 149.7, 147.0, 141.8, 140.2, 135.6, 134.3, 129.8, 129.6, 125.8, 125.1, 122.7, 121.7, 121.4, 120.6, 118.8. MS (ESI) m/z [M + H]^+^ Calculated for C_20_H_16_N_2_O_5_S 397.0858; found 397.0959.

***5-Acrylamido-2-phenoxybenzenesulfonic acid (QB0349)***

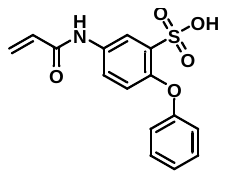

Prepared as shown in *the general experimental procedures B* on a 0.15 mmol scale. Yield = 28% (33 mg); Appearance – White Solid; ^1^H NMR (300 MHz, DMSO-*d*_6_) δ 7.30 – 7.14 (m, 3H), 7.09 – 7.06 (m, 1H), 6.91 (t, *J* = 7.4 Hz, 1H), 6.79 (dt, *J* = 7.8, 1.1 Hz, 3H), 6.56 – 6.42 (m, 3H). ^13^C NMR (75 MHz, DMSO-*d*_6_) δ 166.6, 160.2, 144.8, 141.0, 140.9, 129.3, 122.5, 121.3, 117.8, 115.5, 114.5. MS (ESI) m/z [M + H]^+^ Calculated for C_15_H_13_NO_5_S 320.0593; found 320.0597.

***5-(2-Cyanoacrylamido)-2-phenoxybenzenesulfonic acid (QB0354)***

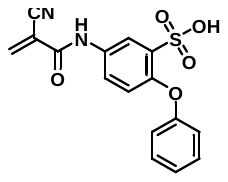

Prepared as shown in *the general experimental procedures B* on a 0.15 mmol scale. Yield = 43% (21 mg); Appearance – White Solid; ^1^H NMR (300 MHz, methanol-*d_4_*) δ 7.38 (s, 1H), 7.30 (dd, *J* = 2.2, 1.1 Hz, 1H), 7.28 – 7.19 (m, 2H), 7.04 – 6.94 (m, 3H), 6.72 – 6.65 (m, 2H). ^13^C NMR (75 MHz, methanol-*d_4_*) δ 165.6, 157.6, 151.8, 139.3, 136.5, 136.4, 130.2, 129.9, 128.4, 124.4, 122.1, 120.6, 117.8, 115.1. MS (ESI) m/z [M + H]^+^ Calculated for C_16_H_12_N_2_O_5_S 345.0545; found 345.0551.

**Synthesis of heteroaryl-sulfonic acids**

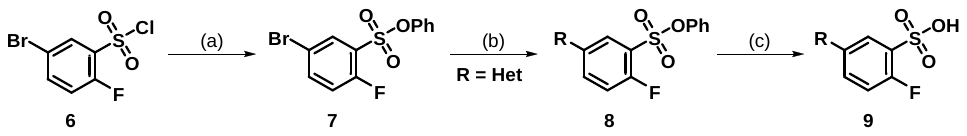

(a) phenol (2.0 equiv.), DIPEA (2.0 equiv.), DCM (0.33 M), rt, 2 h, 65% yield; (b) NH-heteroaryl substrate (1.2 equiv.), Pd(dppf)Cl₂, (10 mol%), Cs_2_CO_3_ (3.0 equiv.), dioxane (0.2 M), 110 ^o^C, 12 h, 24-27% yield; (c) (i) aq. NaOH (2.0 M, 5.0 equiv.), dioxane:water (v/v 3:1, 0.1 M), 100 ^o^C, 6 h. (ii) aq. HCl (2.0 N, pH adjusted to approximately 3~4), 14-28% yield.

***Phenyl 5-bromo-2-fluorobenzenesulfonate (*7**, ***QB0330)***

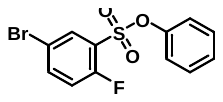

An oven-dried 200 mL single-neck round-bottom flask equipped with a magnetic stir bar was charged with 5-bromo-2-fluorobenzenesulfonyl chloride (**6**) (9.0 g, 33.0 mmol, 1.0 equiv.) and phenol (6.9 g, 66.0 mmol, 2.0 equiv.) in dichloromethane (DCM, 100 mL, 0.33 M). *N,N*-Diisopropylethylamine (DIPEA, 8.5 g, 66.0 mmol, 2.0 equiv.) was then added dropwise to the reaction mixture. The mixture was stirred at room temperature for 2 hours. Upon completion, the reaction was quenched with 50 mL of ice water. The aqueous layer was extracted with DCM (2 × 50 mL), and the combined organic layers were dried over anhydrous sodium sulfate. The solvent was removed under reduced pressure, and the crude product was purified by column chromatography on silica gel (70–230 mesh), using a gradient of ethyl acetate/hexane (5:95 to 10:90 v/v) as the eluent, to afford the desired product, phenyl 5-bromo-2-fluorobenzenesulfonate (**7**, **QB0330**), as a white solid (4.2 g, 65% yield). ^1^H NMR (300 MHz, DMSO-*d*_6_) δ 8.11 (ddd, *J* = 8.9, 4.4, 2.6 Hz, 1H), 7.86 (dd, *J* = 6.1, 2.6 Hz, 1H), 7.63 (dd, *J* = 9.9, 8.9 Hz, 1H), 7.50 – 7.32 (m, 3H), 7.17 – 7.09 (m, 2H).

***C. The general procedure for Pd-catalyzed aryl sulfonate animation***

An oven-dried 50 mL screw-cap reaction vial equipped with a magnetic stir bar was charged with phenyl 5-bromo-2-fluorobenzenesulfonate (**7**, **QB0330**) (1.0 equiv.), the corresponding NH-heteroaryl substrate (1.2 equiv.), dichlorobis(diphenylphosphinoferrocene)palladium(II) (Pd(dppf)Cl₂, 10 mol%), and cesium carbonate (3.0 equiv.) in 1,4-dioxane (0.2 M). The reaction vial was sealed with a screw cap and placed in a pre-heated metal block at 110 °C. The reaction mixture was stirred at 110 °C for 12 hours. After completion, the reaction mixture was cooled to room temperature and quenched with water. The aqueous layer was extracted with ethyl acetate (30 mL × 3) and washed with brine. The combined organic extracts were dried over anhydrous sodium sulfate and concentrated under reduced pressure. The crude product was purified by column chromatography on silica gel using a gradient of dichloromethane/methanol (5:95 to 25:75 v/v) as eluent to afford the desired products (**8**).

***Phenyl 2-fluoro-5-(3-nitro-1H-pyrrol-1-yl)benzenesulfonate (QB0352)***

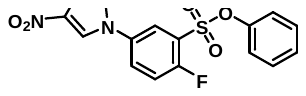

Prepared as shown in *the general experimental procedures C* on a 1.2 mmol scale. Yield = 27% (122 mg); Appearance – White Solid; ^1^H NMR (300 MHz, CDCl_3_) δ 8.06 (d, *J* = 2.5 Hz, 1H), 7.63 (dd, *J* = 8.8, 2.5 Hz, 1H), 7.49 – 7.29 (m, 4H), 7.27 – 7.17 (m, 2H), 7.15 – 7.06 (m, 2H), 6.85 (d, *J* = 8.8 Hz, 1H).

***Phenyl 2-fluoro-5-(4-nitro-2H-1,2,3-triazol-2-yl)benzenesulfonate (QB0357)***

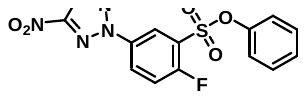

Prepared as shown in *the general experimental procedures C* on a 1.2 mmol scale. Yield = 24% (110 mg); Appearance – White Solid; ^1^H NMR (300 MHz, CDCl_3_) δ 7.94 (d, *J* = 2.5 Hz, 1H), 7.78 (dd, *J* = 8.9, 2.5 Hz, 1H), 7.53 – 7.44 (m, 2H), 7.44 – 7.35 (m, 2H), 7.24 – 7.10 (m, 2H), 6.95 (d, *J* = 8.9 Hz, 1H).

***2-Fluoro-5-(3-nitro-1H-pyrrol-1-yl)benzenesulfonic acid (QB0360)***

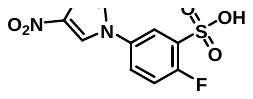

Prepared as shown in *the general experimental procedures B* on a 0.20 mmol scale. Yield = 28% (18 mg); Appearance – White Solid; ^1^H NMR (300 MHz, methanol-*d*_4_) δ 8.07 (d, *J* = 2.5 Hz, 1H), 7.37 (d, *J* = 2.5 Hz, 1H), 7.36 – 7.25 (m, 2H), 7.19 – 6.99 (m, 3H), 6.68 (d, *J* = 8.7 Hz, 1H). ^13^C NMR (75 MHz, methanol-*d*_4_) δ 131.6, 131.6, 129.7, 124.2, 124.1, 120.5, 119.9, 76.6, 49.1. MS (ESI) m/z [M + H]^+^ Calculated for C_10_H_7_FN_2_O_5_S 287.0138; found 287.0137.

***(D) The general procedure for catalytic hydrogenation using Pd/C***

An oven-dried 50 mL round-bottom flask equipped with a magnetic stir bar was charged with the corresponding nitroarene substrate (1.0 equiv.) in dichloromethane (DCM, 0.3 M). Palladium on carbon (Pd/C, 10 wt.% on activated carbon, 10 mol%) was added at room temperature. The flask was evacuated and refilled with a hydrogen balloon. The reaction mixture was stirred at room temperature for 24 hours. After completion of the reaction (monitored by TLC), the mixture was filtered through a bed of Celite. The filtrate was concentrated under reduced pressure, and the crude product was purified by column chromatography on silica gel (70–230 mesh) using a gradient of methanol/dichloromethane (5:95 to 30:70 v/v) as eluent to afford the desired amine derivatives.

***5-(3-Amino-1H-pyrrol-1-yl)-2-phenoxybenzenesulfonic acid (QB0361)***

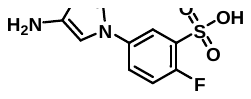

Prepared as shown in *the general experimental procedures D* on a 0.20 mmol scale. Yield = 72% (48 mg); Appearance – White Solid; ^1^H NMR (300 MHz, methanol-*d*_4_) δ 7.97 (dd, *J* = 7.8, 1.7 Hz, 1H), 7.37 (tdd, *J* = 7.3, 4.4, 1.7 Hz, 3H), 7.21 – 7.02 (m, 4H), 6.83 (dd, *J* = 8.2, 1.0 Hz, 1H). ^13^C NMR (75 MHz, methanol-*d*_4_) δ 131.9, 129.4, 128.7, 123.5, 122.0, 119.7, 118.5, 47.4, 47.1, 46.8. MS (ESI) m/z [M + H]^+^ Calculated for C_10_H_9_FN_2_O_3_S 257.0396; found 257.0393.

***5-(4-Nitro-2H-1,2,3-triazol-2-yl)-2-phenoxybenzenesulfonic acid (QB0363)***

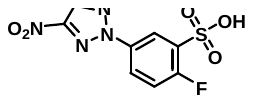

Prepared as shown in *the general experimental procedures B* on a 0.25 mmol scale. Yield = 14% (11 mg); Appearance – White Solid; ^1^H NMR (300 MHz, methanol-*d*_4_) δ 8.13 (d, *J* = 2.5 Hz, 1H), 7.98 (d, *J* = 2.5 Hz, 1H), 7.85 (dd, *J* = 8.7, 2.5 Hz, 1H), 7.82 – 7.68 (m, 2H). ^13^C NMR (75 MHz, methanol-*d*_4_) δ 149.6, 130.1, 129.7, 127.1, 122.1, 119.8, 77.2. MS (ESI) m/z [M + H]^+^ Calculated for C_8_H_5_FN_4_O_5_S 289.0043; found 289.0056.

**Synthesis of** ***3-(Furan-2-yl)-N-(2-(trifluoromethyl)pyridin-4-yl)acrylamide (QB0364)***

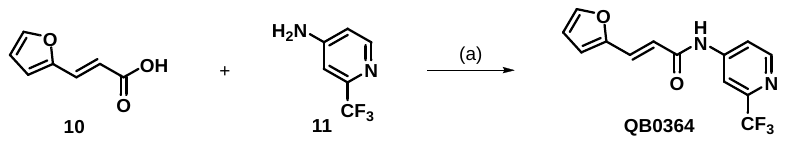

(a) EDC.HCl (2.0 equiv.), DCM (0.1 M), rt, 12 h, 38% yield.

An oven-dried 25 mL two-neck round bottle flask equipped with a magnetic stir bar was charged with 3-(furan-2-yl)acrylic acid (**10**) (82.9 mg, 0.60 mmol, 1.0 equiv.), EDC.HCl (230 mg, 1.2 mmol, 2.0 equiv.) in DCM (6 mL, 0.1 M) solvent. The reaction mixture was stirred for 30 minutes at room temperature, and then 2-(trifluoromethyl)pyridine-4-amine (**11**) (98 mg, 0.60 mmol, 1.0 equiv) was added. The reaction mixture was stirred at room temperature for 12 h. Later, the solvent was evaporated in vacuo and purified by silica gel (70-230 mesh size) column chromatography using methanol/dichloromethane eluent (2:98 to 5:95 v/v) to obtain the desired product as a white solid (**QB0364**), 79 mg in 38% yield. ^1^H NMR (300 MHz, DMSO-*d*_6_) δ 10.54 (s, 1H), 8.31 (d, *J* = 2.7 Hz, 1H), 8.01 (dd, *J* = 9.0, 2.7 Hz, 1H), 7.68 – 7.54 (m, 3H), 7.51 – 7.29 (m, 8H), 7.30 – 7.02 (m, 6H), 6.75 (d, *J* = 15.8 Hz, 1H). ^13^C NMR (75 MHz, DMSO-*d*_6_) δ 164.5, 153.6, 150.9, 150.4, 146.5, 144.9, 130.9, 116.7, 115.9, 115.5, 112.6, 111.1, 106.3, 106.2. MS (ESI) m/z [M + H]^+^ Calculated for C_13_H_9_F_3_N_2_O_2_ 283.0694; found 283.0697.

**Synthesis of *(E)-2-Fluoro-5-(3-(furan-2-yl)acrylamido)benzenesulfonic acid (QB0365)***

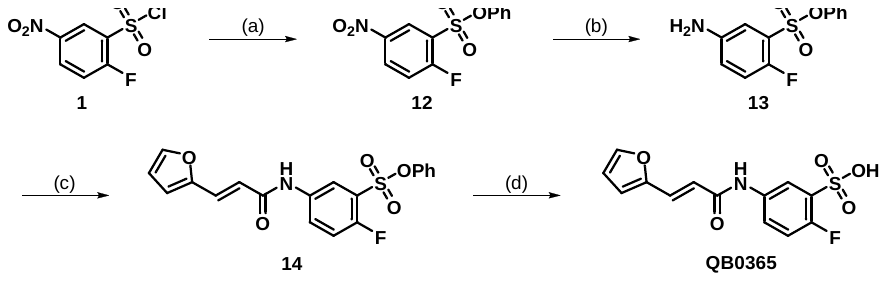

(a) phenol (1.2 equiv.), Et₃N (2.0 equiv.), DCM (0.25 M), 0 ^o^C, 2 h, 92% yield; (b) Pd/C (10 mol%), H_2_, DCM (0.20 M), 25 ^o^C, 24 h, 97% yield; (c) (i) 3-(furan-2-yl)acrylic acid (1.5 equiv.), SOCl_2_ (excess), 80 ^o^C, 2 h; (ii) amine **13** (1.0 equiv.), TEA (2.0 equiv.), DCM (0.2 M), 10 ^o^C, 2 h, 87% yield; (d) (i) aq. NaOH (2.0 M, 5.0 equiv.), dioxane:water (v/v 3:1, 0.1 M), 100 ^o^C, 6 h. (ii) aq. HCl (2.0 N, pH adjusted to approximately 3~4), 62% yield.

***Phenyl 2-fluoro-5-nitrobenzenesulfonate (12, AK0284)***

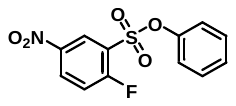

An oven-dried 200 mL single-neck round-bottom flask equipped with a magnetic stir bar was charged with 2-fluoro-5-nitrobenzenesulfonyl chloride (**1**) (4.8 g, 20.0 mmol, 1.0 equiv.) and phenol (2.3 g, 24.0 mmol, 1.2 equiv.) in dichloromethane (DCM, 80 mL, 0.25 M). The reaction mixture was cooled to 0 °C for 30 minutes under an argon atmosphere. Triethylamine (Et₃N, 4.05 g, 40.0 mmol, 2.0 equiv.) was then added dropwise to the reaction mixture using an addition funnel. After the addition was complete, the reaction mixture was stirred at 0 °C for 2 hours. Upon completion, the reaction mixture was quenched with 50 mL of ice water. The aqueous layer was extracted with DCM (3 × 75 mL), and the combined organic extracts were dried over anhydrous sodium sulfate. The solvent was removed under reduced pressure, and the crude product was purified by column chromatography on silica gel (70–230 mesh) using a gradient of ethyl acetate/hexane (5:95 to 10:90 v/v) as eluent to afford the desired product, phenyl 2-fluoro-5-nitrobenzenesulfonate (**12**, **AK0284**), as a yellow oil (5.5 g, 92% yield). ^1^H NMR (300 MHz, CDCl_3_) δ 8.69 (dt, *J* = 8.2, 3.9 Hz, 1H), 8.62 – 8.46 (m, 1H), 7.50 (q, *J* = 5.1, 4.7 Hz, 1H), 7.43 – 7.20 (m, 3H), 7.15 (dd, *J* = 8.3, 4.5 Hz, 2H). ^13^C NMR (75 MHz, CDCl_3_) δ 164.4, 160.8, 149.03, 143.8, 131.9, 131.8, 130.3, 128.1, 127.5, 125.5, 125.3, 121.8, 119.2, 118.8.

***Phenyl 5-amino-2-fluorobenzenesulfonate (13, AK0286)***

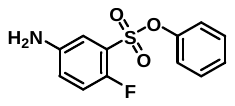

An oven-dried 100 mL single-neck round-bottom flask equipped with a magnetic stir bar was charged with phenyl 2-fluoro-5-nitrobenzenesulfonate (**12**, **AK0284**) (2.97 g, 10.0 mmol, 1.0 equiv.) in dichloromethane (DCM, 50 mL, 0.2 M). Palladium on carbon (Pd/C, 10 wt.% on activated carbon) (106 mg, 10 mol%) was added at room temperature. The flask was evacuated and refilled with a hydrogen balloon. The reaction mixture was stirred at room temperature for 24 hours. After completion of the reaction (monitored by TLC), the mixture was filtered through a bed of Celite. The filtrate was concentrated under reduced pressure, and the crude product was purified by column chromatography on silica gel (70–230 mesh) using a gradient of methanol/dichloromethane (1:99 to 3:97 v/v) as eluent to afford the desired product, phenyl 5-amino-2-fluorobenzenesulfonate (**13**, **AK0286**), as a light-brown solid (2.6 g, 97% yield). ^1^H NMR (300 MHz, CDCl_3_) δ 7.25 (dq, *J* = 14.1, 7.2 Hz, 3H), 7.09 (t, *J* = 6.3 Hz, 2H), 7.02 – 6.89 (m, 2H), 6.83 (dd, *J* = 8.2, 3.9 Hz, 1H), 3.83 (s, 2H). ^13^C NMR (75 MHz, CDCl_3_) δ 153.6, 150.3, 149.4, 143.3, 143.3, 129.8, 127.4, 123.2, 122.9, 122.2, 122.1, 122.0, 118.1, 117.8, 115.7.

***Phenyl (E)-2-fluoro-5-(3-(furan-2-yl)acrylamido)benzenesulfonate (14, AK0297)***

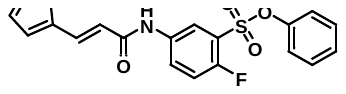

An oven-dried 200 mL screw-cap reaction vial equipped with a magnetic stir bar was charged with 3-(furan-2-yl)acrylic acid (2.2 g, 15.0 mmol, 1.5 equiv.) and excess thionyl chloride (5 mL). The reaction mixture was refluxed at 60 °C for 2 hours. After completion, the excess thionyl chloride was removed under reduced pressure. Dichloromethane (DCM, 30 mL, 0.30 M) was then added, and the resulting reaction mixture was stirred at 0 °C for 15 minutes under an argon atmosphere. Next, a solution of phenyl 5-amino-2-fluorobenzenesulfonate (**13**, **AK0286**) (2.6 g, 10.0 mmol, 1.0 equiv.) and triethylamine (Et₃N, 2.06 g, 20.0 mmol, 2.0 equiv.) in DCM (20 mL) was added dropwise to the reaction mixture using an addition funnel. Upon completion, the reaction was quenched with 50 mL of ice water and extracted with DCM (3 × 50 mL). The combined organic layers were washed with aqueous NaOH (1.0 M, 50 mL), dried over anhydrous sodium sulfate, and concentrated under reduced pressure. The crude product was purified by column chromatography on silica gel (70–230 mesh) using a gradient of ethyl acetate/hexane (15:85 to 25:75 v/v) as eluent to afford the desired product, phenyl (*E*)-2-fluoro-5-(3-(furan-2-yl)acrylamido)benzenesulfonate (**14**, **AK0297**), as a white solid (3.3 g, 87% yield). ^1^H NMR (300 MHz, CDCl_3_) δ 8.23 (dt, *J* = 9.5, 3.5 Hz, 1H), 7.91 (dd, *J* = 6.0, 2.8 Hz, 1H), 7.51 – 7.39 (m, 2H), 7.35 – 7.22 (m, 4H), 7.14 (dd, *J* = 6.1, 3.8 Hz, 2H), 6.62 (d, *J* = 3.6 Hz, 1H), 6.53 – 6.37 (m, 2H). ^13^C NMR (75 MHz, CDCl_3_) δ 165.0, 156.6, 153.2, 151.1, 149.2, 144.4, 135.5, 129.7, 129.1, 127.7, 127.6, 127.4, 123.2, 123.0, 121.8, 121.5, 117.8, 117.7, 117.5, 114.6, 112.2.

***(E)-2-fluoro-5-(3-(furan-2-yl)acrylamido)benzenesulfonic acid (QB0365)***

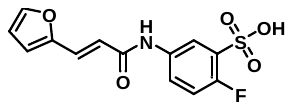

An oven-dried 8 mL screw-cap reaction vial equipped with a magnetic stir bar was charged with (*E*)-2-fluoro-5-(3-(furan-2-yl)acrylamido)benzenesulfonate (**14**, **AK0297**) (78 mg, 0.20 mmol, 1.0 equiv.) and aq. NaOH (2.0 M, 0.5 mL, 5.0 equiv.) in a dioxane:water (v/v 3:1, 2 mL, 0.1 M) mixture. The vial was sealed with a screw cap and placed in a pre-heated metal block at 100 °C for 6 hours. Upon completion of the reaction, the mixture was cooled to room temperature and acidified with 2 N aqueous HCl to adjust the pH to approximately 3~4. The resulting mixture was extracted with ethyl acetate (3 × 10 mL), and the combined organic layers were dried over anhydrous sodium sulfate. The solvent was removed under reduced pressure, and the crude product was purified by column chromatography on silica gel (70–230 mesh), using a gradient of methanol/dichloromethane (10:90 to 25:75 v/v) as the eluent, to afford the desired product, (*E*)-2-fluoro-5-(3-(furan-2-yl)acrylamido)benzenesulfonic acid (**QB0365**), as a white solid (39 mg, 62% yield). ^1^H NMR (300 MHz, CDCl_3_) δ 10.26 (s, 1H), 7.89 (dd, *J* = 6.4, 2.8 Hz, 1H), 7.85 – 7.73 (m, 2H), 7.38 (d, *J* = 15.5 Hz, 1H), 7.07 (t, *J* = 9.2 Hz, 1H), 6.85 (d, *J* = 3.4 Hz, 1H), 6.64 – 6.53 (m, 2H). ^13^C NMR (75 MHz, CDCl_3_) δ 163.7, 156.4, 153.2, 151.4, 145.6, 136.1, 135.8, 134.9, 134.9, 127.7, 121.5, 121.4, 120.3, 120.2, 119.7, 116.6, 116.3, 115.0, 113.0. MS (ESI) m/z [M + H]^+^ Calculated for C_13_H_10_FNO_5_S 312.0342; found 312.0340.

**Supplemental Figures:**

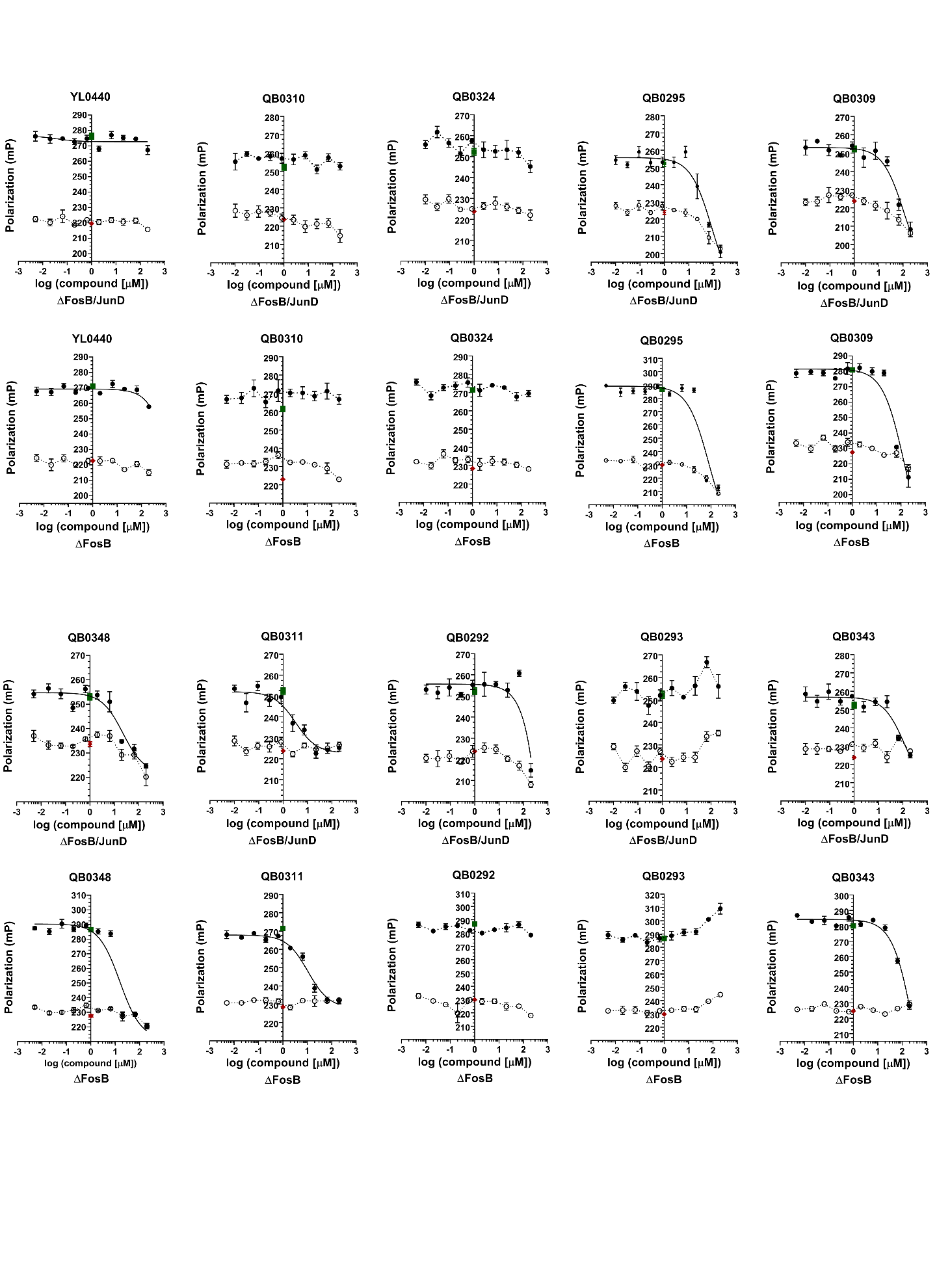

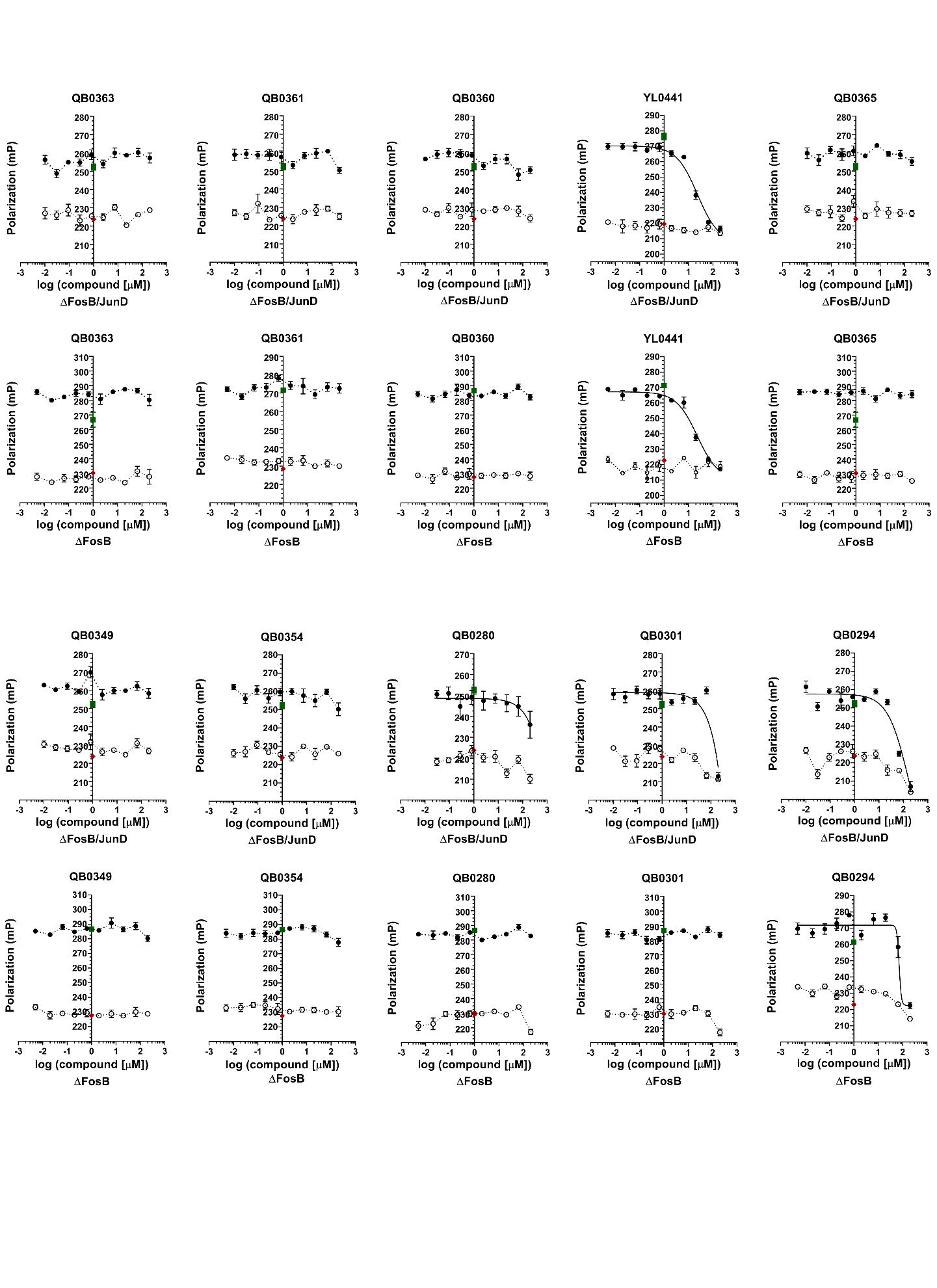

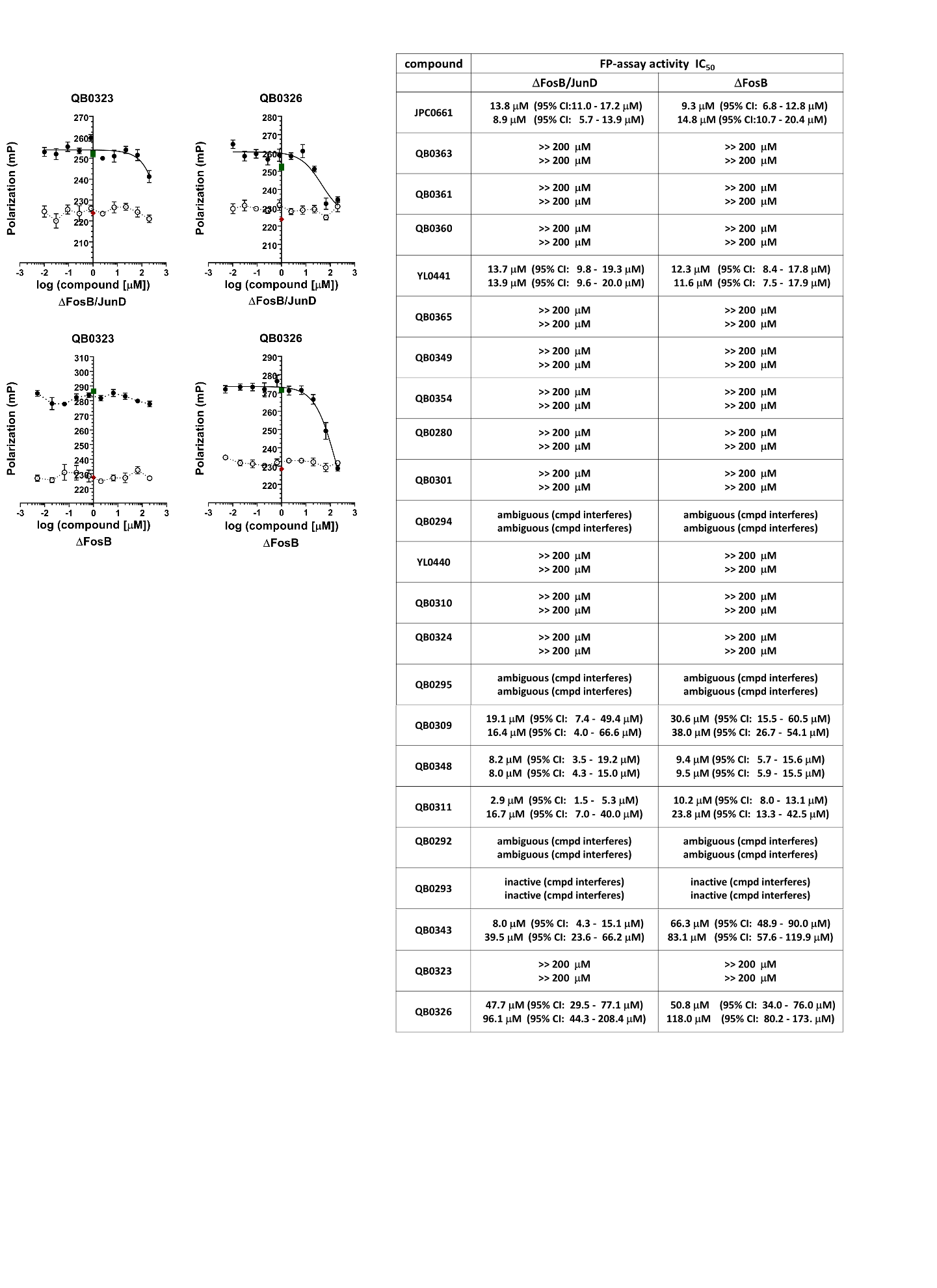

**Figure. S1. Ability of 22 analogues of JPC0661 to disrupt DNA-binding using an FP-DRC assay. A)** Representative FP-DRCs for 22 analogues of JPC0661. The identical FP-DRC plots for YL0441, QB0309, QB0348, QB0311, QB0343 and QB0326 are shown in **Fig. 4** in the main text as well. See **Fig. 4** for an extensive description. **B)** Summary of the compound activities from two independent FP-DRC assays (IC_50_) carried out for the 22 analogues. Compounds displaying an inhibitory activity on protein:DNA binding, but for which the compound also impacted the FP signal of TMR-cdk5 oligo alone are listed as ‘ambiguous (cmpd interferes)’. Compounds with no apparent effect on protein:DNA binding, but which impacted the FP signal of TMR-cdk5 oligo alone are listed as ‘inactive (cmpd interferes)’.

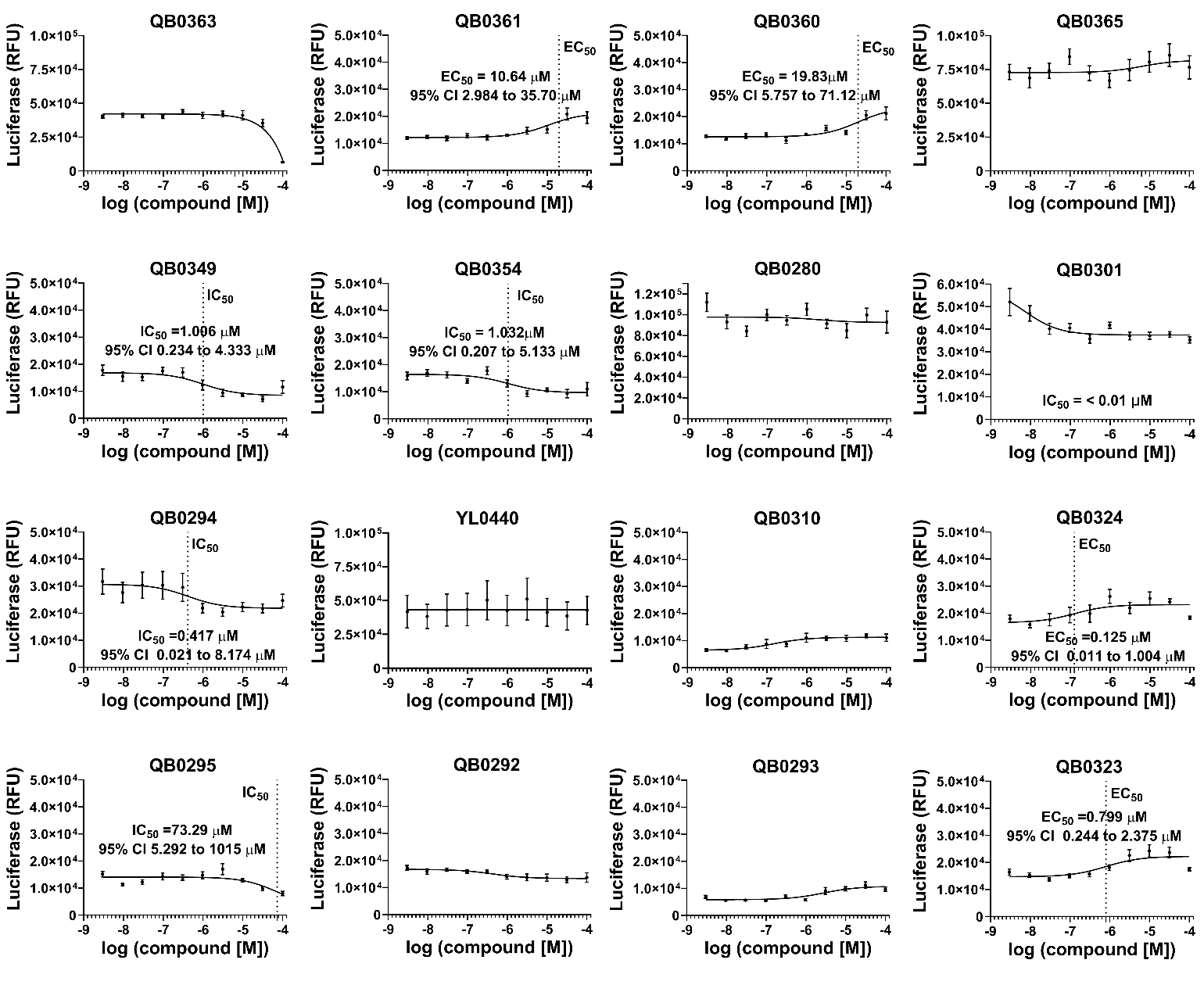

**Figure. S2. Additional analogues of JPC0661 analogues tested in an AP1-reporter assay.** Effects of compounds (0–100 μM) on AP1-driven luciferase activity as evaluated in AP1-luc HEK293 cells. Dose-dependent activation of the AP1 reporter was quantified by measuring changes in luciferase signal, expressed as relative fluorescence units (RFU). Each compound was tested in at least two independent experiments (n = 4 wells per experiment), resulting typically in at least n=~6–8 wells per concentration of compound, and the results were then normalized to the luciferase signal from blank wells within each experiment. Data were fitted to a three-parameter logistic model to determine IC_50_ values and associated 95% confidence intervals (CI). Data points are presented as mean ± SEM.

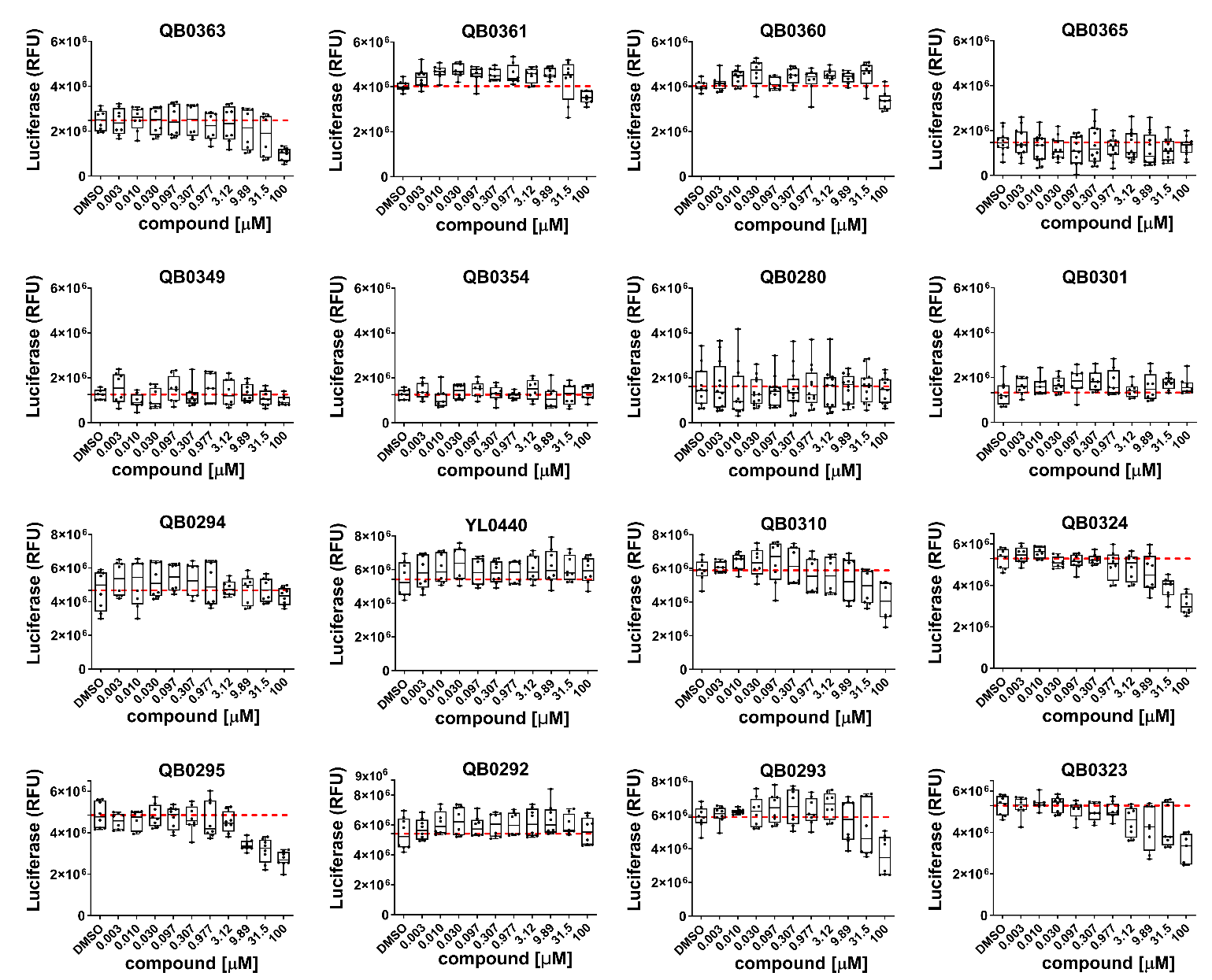

**Figure. S3. Additional analogues of JPC0661 analogues tested in a cell viability assay.** The effect of serially diluted compounds (0.003–100 µM) on the viability of AP1-luc HEK293 cells was assessed using the CellTiter-Glo viability assay after 2 hours. Cell viability was normalized to the control containing 0.5% (v/v) DMSO in the absence of compound (n = 7 wells). Data points represent the mean ± SEM (n=10-12 wells analyzed per concentration).

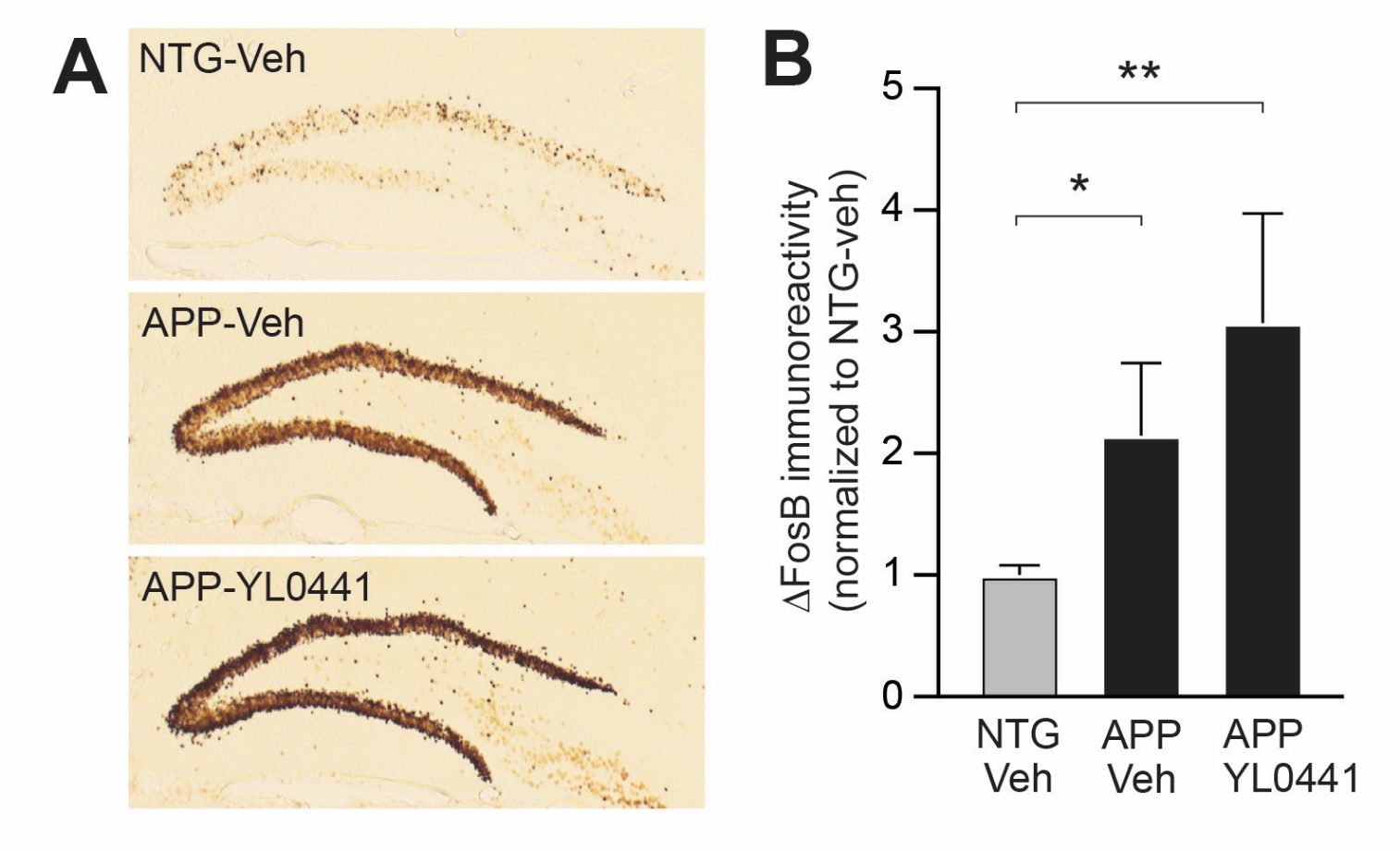

**Supplemental Figure S4. Expression of ΔFOSB in the dentate gyrus of mice treated with vehicle or YL0441. A)** Representative images showing immunohistochemical staining of ΔFOSB from the hemibrain contralateral to the side that received vehicle/YL0441 infusion into the hippocampus in nontransgenic (NTG) or APP mice. **B)** Quantification of ΔFOSB expression in the dentate granule cell layer of NTG or APP mice infused with vehicle or YL0441. n = 5 (NTG), 10 (APP-veh), 11 (APP-LY0441). Kruskal-Wallis test revealed a significant effect of group (p = 0.025); Benjamini Hochberg correction for multiple comparison tests indicated that NTG-veh differed from APP-Veh (p = 0.033) and from APP-YL0441 (p = 0.0074). *p < 0.05, **p < 0.01.

| **compound** | **Fluorescence Polarization-DRC**  **IC_50_ [μM]** | | **AP1-luciferase Reporter Assay**  **IC_50_ [μM]** | **Cell Viability Assay**  **[decrease at 31.5 μM]** |
| --- | --- | --- | --- | --- |
|  | **ΔFOSB/**  **JUND** | **ΔFOSB/**  **ΔFOSB** |  |  |
| **JPC0661** | **+++** | **+++** | **++** | **no** |
| **QB0363** | **-** | **-** | **-** | **yes** |
| **QB0361** | **-** | **-** | **-** | **no** |
| **QB0360** | **-** | **-** | **-** | **no** |
| **YL0441** | **+++** | **+++** | **+++** | **no** |
| **QB0365** | **-** | **-** | **-** | **yes** |
| **QB0349** | **-** | **-** | **++** | **yes** |
| **QB0354** | **-** | **-** | **++** | **no** |
| **QB0280** | **-** | **-** | **-** | **no** |
| **QB0301** | **-** | **-** | **+++** | **no** |
| **QB0294** | **ambiguous** | **ambiguous** | **++** | **no** |
| **YL0440** | **-** | **-** | **-** | **no** |
| **QB0310** | **-** | **-** | **-** | **yes** |
| **QB0324** | **-** | **-** | **-** | **yes** |
| **QB0295** | **ambiguous** | **ambiguous** | **-** | **yes** |
| **QB0309** | **+++** | **++** | **++** | **yes** |
| **QB0348** | **+++** | **+++** | **++** | **yes** |
| **QB0311** | **+++** | **+++** | **-** | **yes** |
| **QB0292** | **ambiguous** | **ambiguous** | **-** | **no** |
| **QB0293** | **ambiguous/inactive** | **ambiguous/inactive** | **-** | **yes** |
| **QB0343** | **++** | **+** | **-** | **no** |
| **QB0323** | **-** | **-** | **-** | **yes** |
| **QB0326** | **+** | **+** | **-** | **no** |

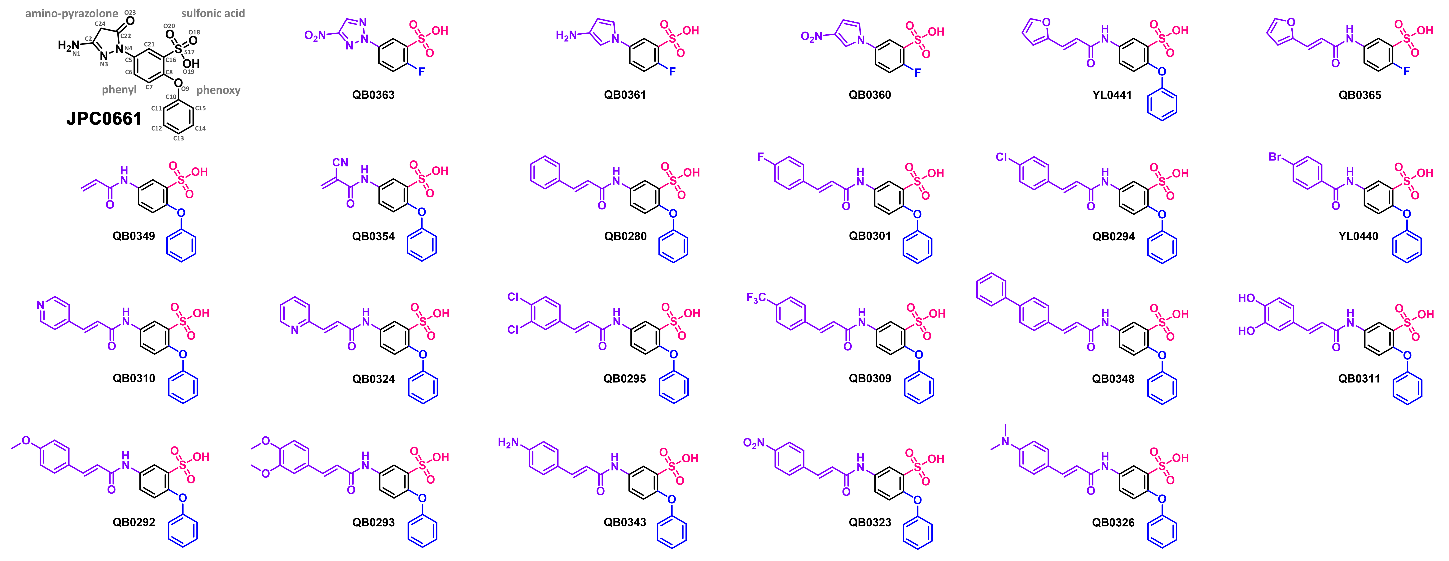

| **activity** | ***FP-assay***  ***IC_50_ (*μM*)*** | ***AP1-Luc reporter assay***  ***IC_50_ (*μM*)*** |
| --- | --- | --- |
| **+++** | **< 20** | **≤ 0.1** |
| **++** | **20 - 50** | **0.1 - 1.0** |
| **+** | **50 - 200** | **1.0 - 10.0** |
| **-** | **> 200** | **> 10.0** |

**Table S1: Medicinal chemistry campaign to optimize lead compound, JPC0661.** Summary of the biochemical and cell-based assays used to rank order 22 analogues of our lead compound JPC0661.
